# Cell fusion after wounding requires plasma membrane damage and endocytosis

**DOI:** 10.64898/2026.05.31.728998

**Authors:** Junmin Hua, Evan S. Krystofiak, Andrew D. Pumford, Andrea Page-McCaw, M. Shane Hutson

## Abstract

Tissue wounds comprise both dead and damaged cells. In epithelial wounds, repair is accomplished by cells at the wound edges, which are themselves often damaged. In the *Drosophila* pupal notum, wound-adjacent epithelial cells with plasma membrane damage often fuse to form syncytia; when plasma membrane damage is prevented, syncytia do not form. Damaged cells share cytoplasm as soon as milliseconds after wounding, and fusion pores connecting cell membranes form minutes later. A genetic screen reveals that wound-induced fusion requires endocytosis machinery, and dynamin localization indicates that endocytosis preferentially targets plasma membrane removed during fusion. Endocytosis promotes cell fusion by specifically promoting fusion pore expansion, indicated by quantitative analysis of cytoplasmic sharing between cells over time. Without endocytosis-mediated cell fusion, wound healing is slowed. Together, our results support a model of damage-induced cell fusion in which plasma membrane damage initiates fusion pores and endocytosis expands fusion pores, resulting in cellular fusion as an integration of single cell damage with tissue repair.

## Introduction

Tissue repair is critical for organismal survival, and understanding how wounds are repaired has both biological and clinical significance. At the simplest level, an epithelial wound can be described as a collection of dead and damaged cells in an otherwise healthy cell sheet. Importantly, some of the damaged cells die while others survive, and the surviving damaged cells are at the front line of healing the wound. These cells actively contribute to tissue repair while simultaneously managing their own cellular damage. How do these damaged cells integrate cell- and tissue-level repair to survive and reconstitute epithelial barrier function? Previously, we reported that wounds in the *Drosophila* notum are repaired by multinucleated syncytia that form specifically in response to injury through multiple rounds of cell-cell fusion. These syncytia rapidly crawl and stretch into the wounded area to reseal the epithelium^1^, but little is known about the triggers and mechanisms leading to wound-induced cell fusion. Here, we report that wound-induced fusion is initiated by cell-level damage to the plasma membrane and that complete fusion requires endocytosis; using these findings we develop a model of wound-induced cell fusion. Our experimental system combines laser-wounding, which generates reproducible and radially symmetric gradients of cellular damage, with live imaging and conditional genetic manipulations in the *Drosophila* pupal notum, to analyze how the integration of cell- and tissue-level wound responses result in cell fusion.

Cell-cell fusion is critical for the repair of several tissues including muscles^2–4^, liver^5^, central nervous system^6,7^, and epidermis^1,8^. It has important applications in the clinic, as transplanted bone marrow cells facilitate repair and regeneration by fusing with damaged resident myocytes^9–17^, hepatocytes^18–24^, Purkinje neurons^24–28^, and Müller glial cells^29,30^. Despite the importance and prevalence of wound-induced cell fusion, its mechanisms are poorly understood. Previous work has identified several genes that contribute to wound-induced fusion, including Rac^8^, Hippo/Yki^31,32^, autophagy genes^33^, β-integrin^31,34^, JNK and JAK/STAT^35^. Nevertheless, the respective role of each component in the fusion process remains largely unclear. Indeed, there has been no overarching model to explain wound-induced fusion.

The better-understood process of developmentally regulated cell fusion may provide a useful framework. Developmental fusion is described by a pore-expansion model (also known as the stalk-pore model), which roughly divides developmental fusion into two steps: (1) fusion pore formation, in which a nanometer-scale pore connects the plasma membranes of two independent cells and makes their cytoplasm continuous; and (2) fusion pore expansion, in which the pore is stabilized and enlarged to micron-scale, eventually eliminating any morphological distinction between the cells^36^. Fusion pore formation requires overcoming the repulsive electrostatic forces that normally keep plasma membranes over 10 nm apart^37^. This energy barrier is overcome by fusogenic proteins or fusogens, which bring opposing plasma membranes within 2 nm of one another^38,39^, for example syncytins during trophoblast fusion^40–42^, myomaker and myomerger during muscle fusion^43–47^, and EFF/AFF during nematode epithelial fusion^48^. Subsequent expansion of a fusion pore is also an energy demanding process. Both modeling^49,50^ and experiments on protein-free lipid bilayers^51^ have demonstrated that fusion pore expansion requires high membrane tension. Applying this framework to wound-induced cell fusions raises a key question: how does damage induce cells to fuse when they have no readily available fusion machinery?

Here we show that wound-induced cell fusion is primed by plasma membrane damage that occurs concomitant with wounding. Cytoplasmic transfer can begin within seconds, and the first fusion pores are established minutes later, but plasma membrane damage and fusion pore formation are not sufficient to drive complete cell-cell fusion. In a small screen of candidate genes, we discovered that cell fusion around wounds requires genes involved in endocytosis.

Quantitative analysis of cytoplasmic transfer during cell fusion demonstrates that wound-induced fusion pores are unstable, expanding and contracting, and mutant analysis shows that endocytosis promotes fusion pore expansion. Thus, both physical damage and cellular processes are required to drive wound-induced cell fusion, resulting in syncytia that then rapidly reseal the epithelium^1^ – a striking integration of damage repair at cellular and tissue levels.

## Results

### Wound-induced fusion is spatially correlated with plasma membrane damage

The *Drosophila* pupal notum is a monolayer columnar epithelium, comprising diploid, mitotically active cells. Although non-fusogenic in healthy pupae, these cells fuse in response to tissue damage (Fig. 1A-C). To understand how, we wounded the living notal epithelium with a single-pulse laser, which inflicts a gradient of cellular damage in concentric rings: for example, we previously reported that the central disc-shaped zone of cell lysis is surrounded by a more distal ring of plasma membrane damage^52^. Plasma membrane damage can be imaged milliseconds after wounding by the immediate (millisecond) influx of extracellular calcium^53^, which is ∼10,000X more concentrated outside than inside the cell (Fig. 1D). We used spinning disk confocal microscopy and the calcium reporter GCaMP6m to image plasma membrane damage, simultaneously imaging cell borders labeled by the adherens junction protein p120ctn-RFP (Fig. 1D’). Within 30 ms of wounding, cells near the wound center were heavily damaged and filled with Ca^2+^. Yet further out, in an annulus of ∼60-90 µm from the wound center, the GCaMP signal revealed discrete sites of membrane damage, appearing as short stripes encircling the wound (Fig. 1D). Their decreasing signal intensity further from to the wound indicated fewer sites of calcium entry and less damage. These GCaMP stripes colocalized perfectly with cell borders, labeled by p120ctn-RFP (Fig. 1D’,E). Hence, plasma membrane damage is concentrated along circumferential cell-cell borders and decreases with distance from the wound center, consistent with the pattern of shear stresses experienced by cells during laser-wounding^54^.

**Figure 1:**
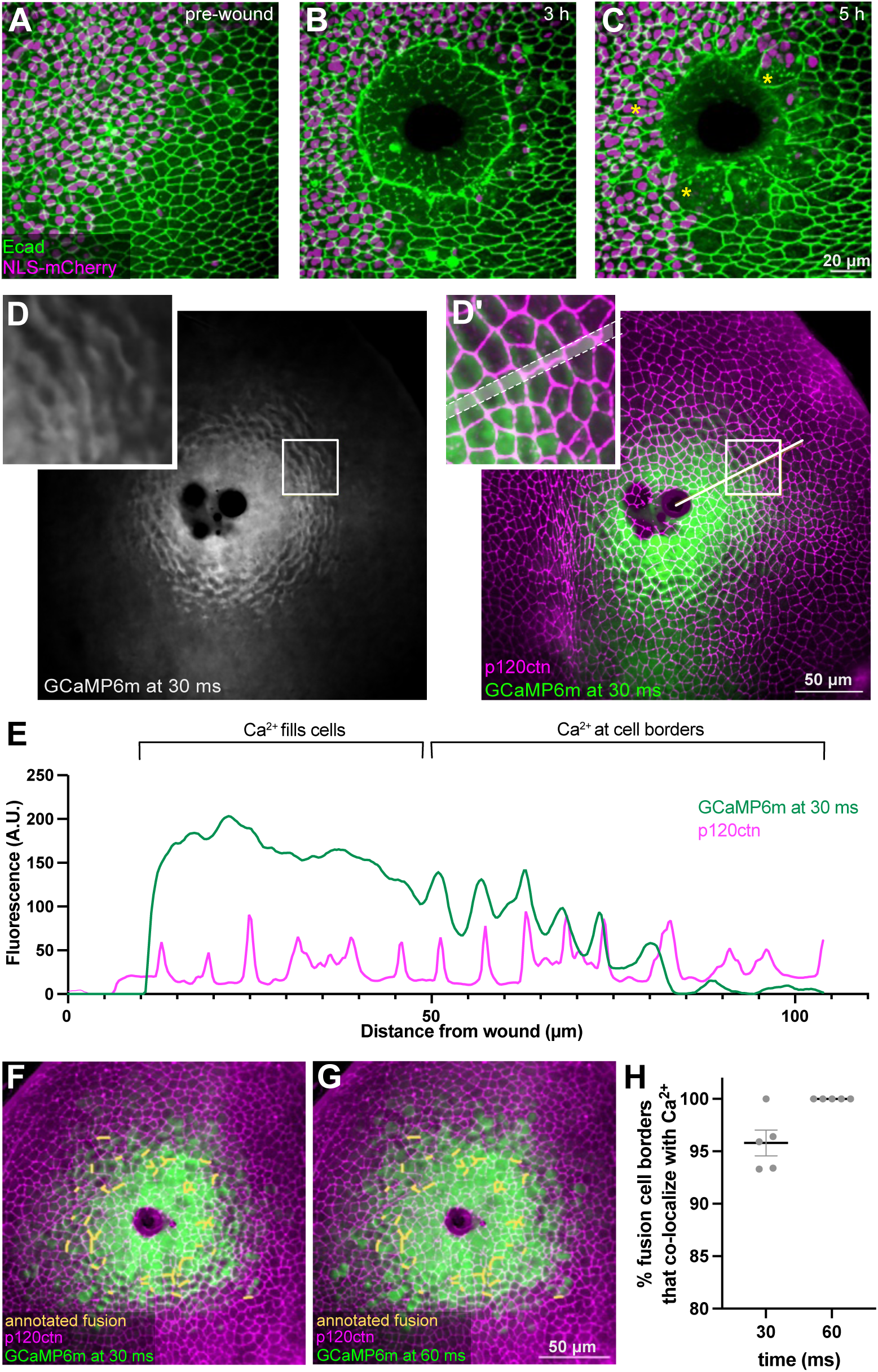
After wounding, cells fuse at sites of plasma membrane damage. **A-C.** Time-lapse images showing fusion of diploid cells into multinucleated syncytia. Asterisks indicate three syncytia; many more are evident. Ecad-GFP labels cell borders; NLS-mCherry labels nuclei in half the wound, in the *pnr* domain. **D-D’.** Sites of plasma membrane damage around a wound visualized by the influx of extracellular calcium, imaged with GCaMP6m at 30 ms after wounding. Plasma membrane damage is concentrated at cell-cell borders, labeled by p120ctn-RFP imaged 10 sec after wounding. Profile plot taken along the white line in D’ shows alignment of damage with cell-cell borders. **F-G.** Yellow lines annotate sites where fusion was observed within 1.5 h of wounding. These sites co-localize with earlier sites of rapid calcium influx (green) at 30 ms (F) or 60 ms (G) after wounding. **H.** All sites of fusion correspond to sites of plasma membrane damage evident within 60 ms of wounding.

We analyzed the spatial relationship between plasma membrane damage and later cell-cell fusion events, which were tracked by the loss of p120ctn between fusing cells. Although this loss is a late indicator of cell fusion, it reliably identifies cell pairs that have already started to exchange cytoplasm^1^. We refer to cell borders that lose the p120ctn adherens junction marker as “fusion borders”. For pupae wounded 15-18 h after puparium formation (APF), every fusion border was located at a previously damaged cell-cell border (n = 187 fusion borders): in 96% of cases, membrane damage at the fusion border was evident via GCaMP fluorescence within 30 ms after wounding; in the remaining 4%, located more distal to the wound, damage became evident within 60 ms (Fig. 1F-H). Thus, cell fusion occurred at sites of wound-induced plasma membrane damage.

### Plasma membrane damage is necessary for wound-induced fusion

Given their spatial correlation, we asked whether plasma membrane damage is required for wound-induced fusion. To make wounds without plasma membrane damage in the surrounding surviving cells, we modified our wounding method. Our standard laser-wounds are generated by a single pulse from a nanosecond UV laser, which initiates a plasma-mediated process that drives explosive vaporization and forms a ∼200-µm diameter cavitation bubble that expands and collapses within microseconds. The rapid movements of the bubble wall produce shear stresses that lyse cells in the wound center and cause collateral membrane damage in surrounding cells^53,55^ (Fig. 2A). To try to eliminate plasma membrane damage, we drastically reduced cavitation bubble size by administering a series of much lower-energy laser pulses^55^ that were repeatedly scanned across the tissue to generate an area comparable to our standard wound area (Fig. 2B)^56^. Killing all targeted cells required rescanning the area three times, taking several minutes. This amount of time impeded our use of Ca^2+^ influx to image plasma membrane damage because Ca^2+^ is released from intracellular stores within ∼45 s after wounding^57^. Thus, to visualize plasma membrane damage, we used the genetically-encoded voltage indicator ArcLight^58,59^, which loses fluorescence when an epithelial cell’s plasma membrane is damaged and thus depolarizes^53^. For our standard single-pulse laser wounds, ArcLight indicated that a wide swath of cells around the wound depolarized, as expected (Fig. 2C-C’’)^53^. For scanned wounds of similar size, surrounding cells maintained their polarized membrane potential (Fig. 2D-D’’), confirming that scanning ablation creates wounds without damaging surrounding cells’ plasma membranes.

**Figure 2:**
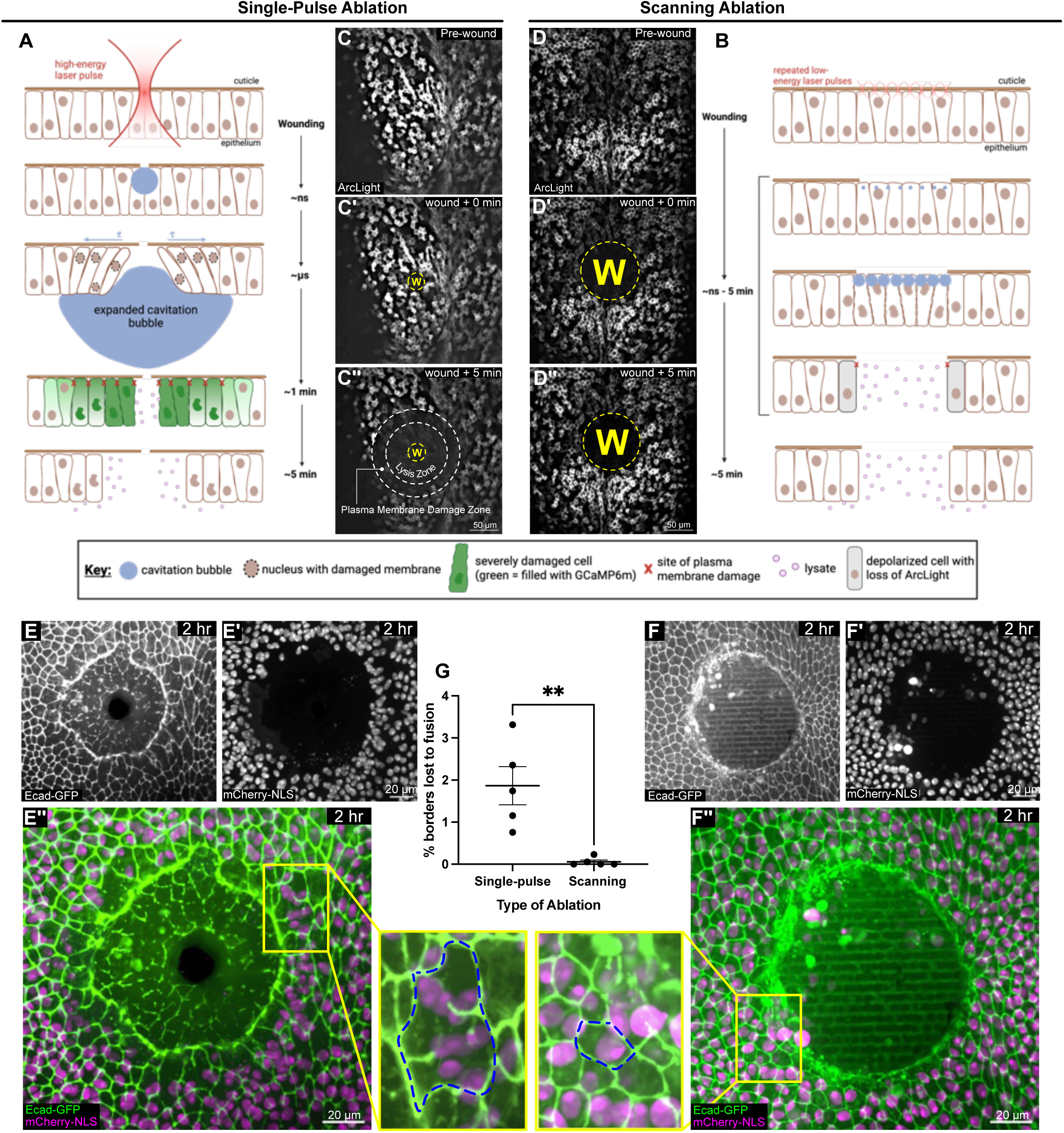
Plasma membrane damage is required for wound-induced fusion. **A-B.** Schematics of two laser wounding protocols applied to the notum epithelium: single-pulse ablation (A) and scanning ablation (B). **C-C’’.** For single-pulse ablation, the central zone of cell lysis is surrounded by a ring of membrane depolarization, indicating a region with plasma membrane damage. Lysis zone identified by later loss of p120ctn-RFP signal; depolarization imaged with ArcLight. **D-D’’.** For scanning ablation, cells are killed throughout a scanned region matching the zone of lysis for single-pulse ablation (C"), but without any surrounding region of membrane depolarization, indicating minimal-to-no plasma membrane damage. **E-E’’**. After single-pulse ablation, cell fuse to form multinucleated syncytia. **F-F’’.** After scanning ablation, virtually no cell-cell fusion is observed. **G.** Comparison of the frequency of fusion after single-pulse and scanning ablation: n = 5 wounds for each protocol; p = 0.0041, unpaired T-test; error bars denote SEM.

Confident that plasma membrane damage is nearly eliminated around scanning-ablation wounds, we compared the frequency of cell fusion events around scanned and standard single-pulse wounds. Importantly, when plasma membrane damage was eliminated, subsequent cell fusion was also eliminated (Fig. 2E-G). We conclude that wound-induced cell fusion requires plasma membrane damage.

### The onset of cell fusion

If plasma membrane damage is fusogenic, we would expect that cell fusion would be initiated soon after wounding. As we previously reported, although the breakdown of adherens junctions between fusing cells occurs predominantly within an hour after injury, fusing cells can start sharing cytoplasm more than 10 minutes earlier^1^. To better understand the temporal dynamics of cytoplasmic sharing, we tracked the transfer of cytoplasmic GFP from clonally labeled donor cells, generated by a conditional flip-out Gal4 cassette, to unlabeled recipient cells. As shown in Fig. 3A, we observed multiple donor-receptor fusion partners around a single wound and noted that fusion partners initiate cytoplasmic sharing at different times, ranging from milliseconds to hours after wounding. When imaged at high temporal resolution (500-ms intervals), some recipient cells increased fluorescence immediately after wounding (Fig. 3B,C), with fluorescence levels well above those of non-recipient bystander cells (Fig. 3B-D). Among ten donor-recipient-bystander triplets, four initiated wound-induced transfer of GFP specifically to the recipient within 500 ms of injury (Fig. 3E, Fig. S1A-C); the remaining six initiated GFP transfer specifically to the recipient at later times, ranging from 5 seconds to over 100 seconds after wounding (Fig. 3E, Fig. S1D-H). These results are consistent with fusion being triggered at wounding by plasma membrane damage.

**Figure 3:**
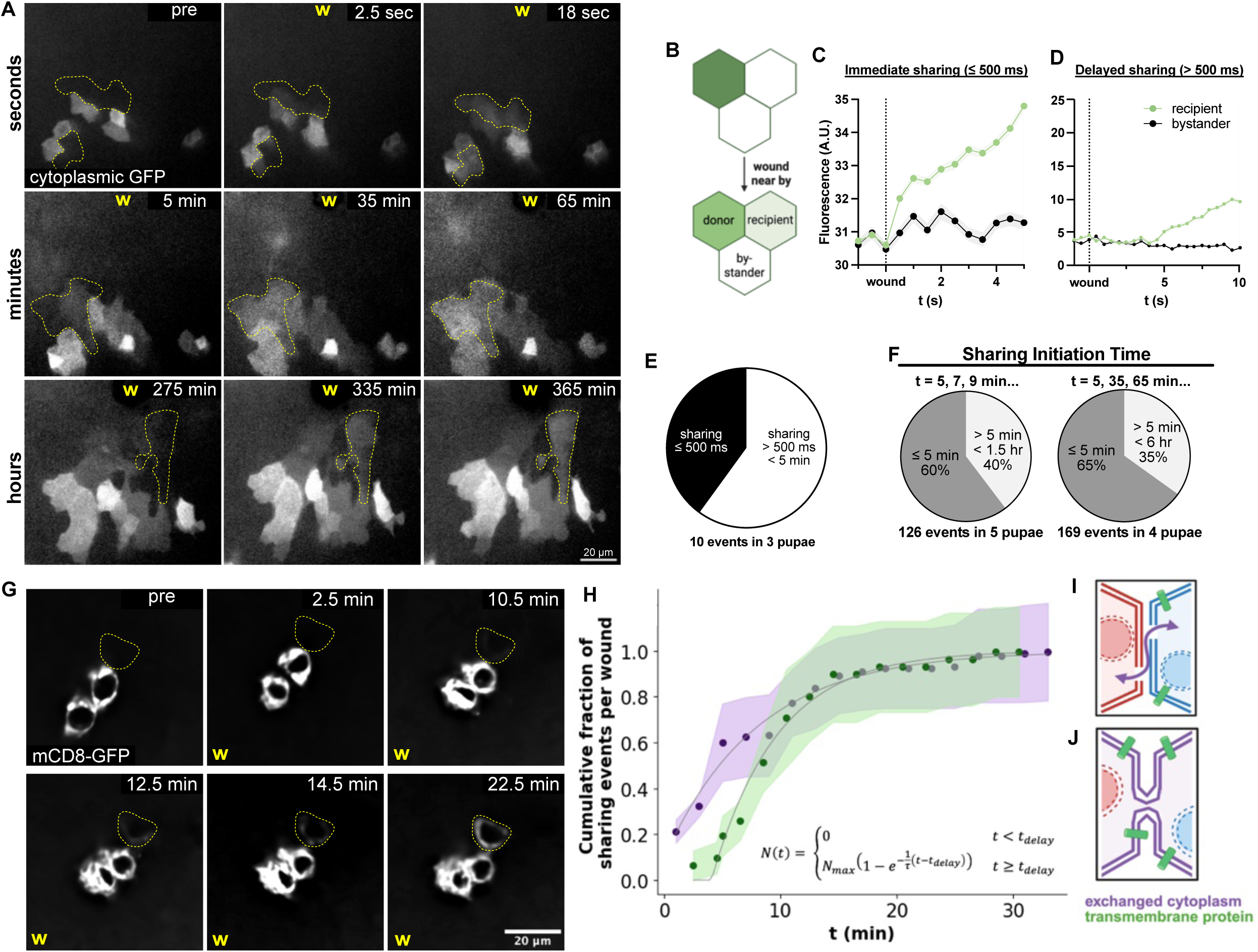
Wound-induced cell fusion initiates over a broad time range, from milliseconds to hours after wounding. **A.** Time-lapse images collected after a single wound. Cell fusion is identified by the transfer of cytoplasmic GFP from a labeled donor cell to an unlabeled recipient, occurring at seconds, minutes, or hours after wounding as shown. Recipients are outlined by yellow dotted lines. **B-E.** By measuring cytoplasmic GFP levels over time in donor cells, recipient cells, and adjacent bystander cells (B), fusion initiation times were determined at half-second resolution for 10 cell fusion events. GFP transfer can occur within 500 ms after wounding (C), or at later times (D). In 4/10 events, fusion was apparent within 500 ms after wounding; in the other 6/10, fusion was delayed by up to 100 seconds (E). **C.** Analyses of GFP transfer over longer time scales – with temporal resolution of 2 min or 30 min – show that about one-third of cytoplasmic GFP sharing events initiated more than 5 min after wounding. **D.** Time-lapse images collected after a single wound, with fusion identified by the initiation of transfer/sharing of membrane-bound mCD8-GFP. A single z-slice is shown. **E.** Cumulative fraction of cytoplasmic (green) and membrane-bound (magenta) GFP transfers initiated over time. Cytoplasmic sharing initiates earlier, but the two curves are highly overlapped for times longer than 10 min. Fitting these data to exponential approaches to saturation (equation on graph) shows that cytoplasmic transfer was initiated with a delay of -0.8 +/- 0.9 min and a time-constant of 8.0 +/- 1.6 min (95% confidence limits; n = 5 wounds), while membrane-bound transfer was initiated with a longer delay, 4.1 +/- 0.4 min, but a shorter time constant, 5.6 +/- 1.1 min (n =11 wounds). Data collected at 2-min intervals; shading indicates SEM. **I-J.** Cartoons illustrate that cytoplasmic sharing can take place via open channels between cells (I), but membrane sharing requires the formation of a membrane-lined fusion pore (J).

When imaged for longer periods after wounding (at 2- or 30-min intervals), two-thirds of cytoplasmic GFP transfer events took place within 5 min of wounding, but the remaining third initiated minutes to hours later (Fig. 3F). The cumulative count of fusion initiations over time was fit well by an exponential approach to saturation (Fig. 3H, green curve)

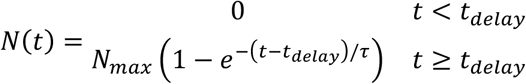

where sharings begin with minimal delay, *t_delay_* = -0.8 ± 0.9 min, and accumulate with a time constant 1″ = 8.0 ± 1.6 min (Fig. 3H). The timescale of minutes is surprising, given previous studies that reported provisional plasma membrane repair within just ∼30 sec^60^. The longer time constant indicates that the fusion of damaged membranes is not simply caused by two cells cross-sealing short-lived free edges of their broken plasma membranes. It instead suggests that wound-induced fusion is a regulated process.

To investigate further, we visualized the transfer of a fluorescent transmembrane protein, mCD8-GFP, from clonally labeled donor cell membranes to recipient cell membranes (Fig. 3G). Since sharing of mCD8-GFP requires a continuous plasma membrane between the fusion partners, it is a direct indicator of fusion pore formation. In contrast with cytoplasmic GFP, sharing of mCD8-GFP began only after a substantial delay, *t_delay_* = 4.1 ± 0.4 min (Fig. 3H); these sharing events then accumulated with a time constant τ = 5.6 ± 1.1 min. The delay in sharing of membrane-bound GFP is important. First, it tells us that the very early sharing of cytoplasmic GFP occurs through open rather than membrane-lined channels (Fig. 3I). Second, because provisional plasma membrane repair takes seconds-to-minutes^60^, it suggests that damaged plasma membranes are provisionally repaired before fusion pores form (Fig. 3J). Indeed, the idea that provisionally repaired membranes are fusogenic can explain the timescale of fusion, which occurs minutes-to-hours after wounding (Fig. 3A).

To confirm the existence of fusion pores, we used TEM to image epithelial cells of the pupal notum after multiple laser ablations along the dorsal midline. We observed fusion pores in multiple biological samples. In each case, membrane bilayers of two cells were continuous, with cytoplasmic material like ribosomes filling the connected space between neighboring nuclei, suggesting that these are true pores between cells (Fig. 4). Interestingly, wound-induced fusion events were often associated with multiple fusion pores of various sizes – quite different from the one fusion pore per fusion event observed for developmentally regulated fusion in *Drosophila* myogenesis^61^. The difference is likely that wound-induced fusion is primed by provisionally-repaired plasma membrane damage present throughout the membrane.

**Figure 4:**
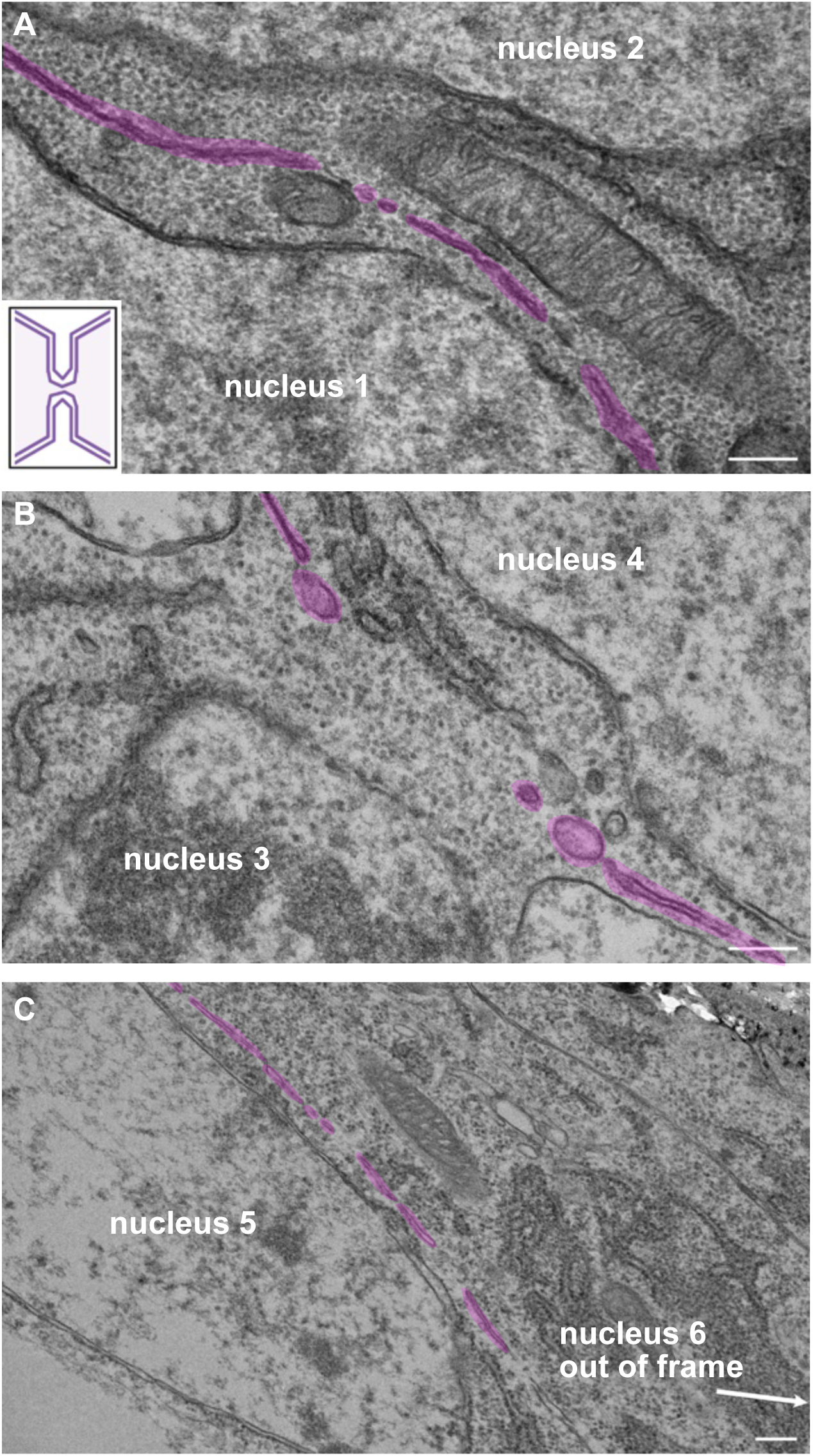
Multiple fusion pores of various sizes can be observed after wounding. **A-C.** Three example TEM images showing the presence of fusion pores in samples fixed 30 min after wounding. Fusion pores are visible as interruptions in the double plasma membranes separating adjacent cells, highlighted in magenta. Scale bars = 200 nm.

Altogether, our data indicate that immediately after wounding plasma membrane damage creates open channels that are transformed into fusion pores at multiple locations via provisionally-repaired plasma membranes.

### A candidate screen identifies endocytosis genes required for cell fusion after wounding

To learn more, we conducted a candidate screen to identify regulators of wound-induced fusion. We knocked down or inhibited gene products in a temporally and spatially restricted manner with *Gal80^ts^*limiting expression to the pupal stage of development and *pnr-Gal4* limiting expression to only one half of the wound bed, with the other half serving as an internal control^57^. Laser wounds were targeted to either the left or right edge of the *pnr* domain, allowing comparison of normal cells on the control side to loss-of-function cells labeled with NLS-mCherry on the *pnr* side to (Figs. 1A, 5A), and we assayed the frequency of fusion by the loss of adherens junctions (Ecadherin-GFP) over time accompanied by the appearance of large syncytial cells. Before screening, we confirmed that fusion was comparable in *pnr* and control domains when there was no functional genetic manipulation (Fig. 5B, B’).

**Figure 5:**
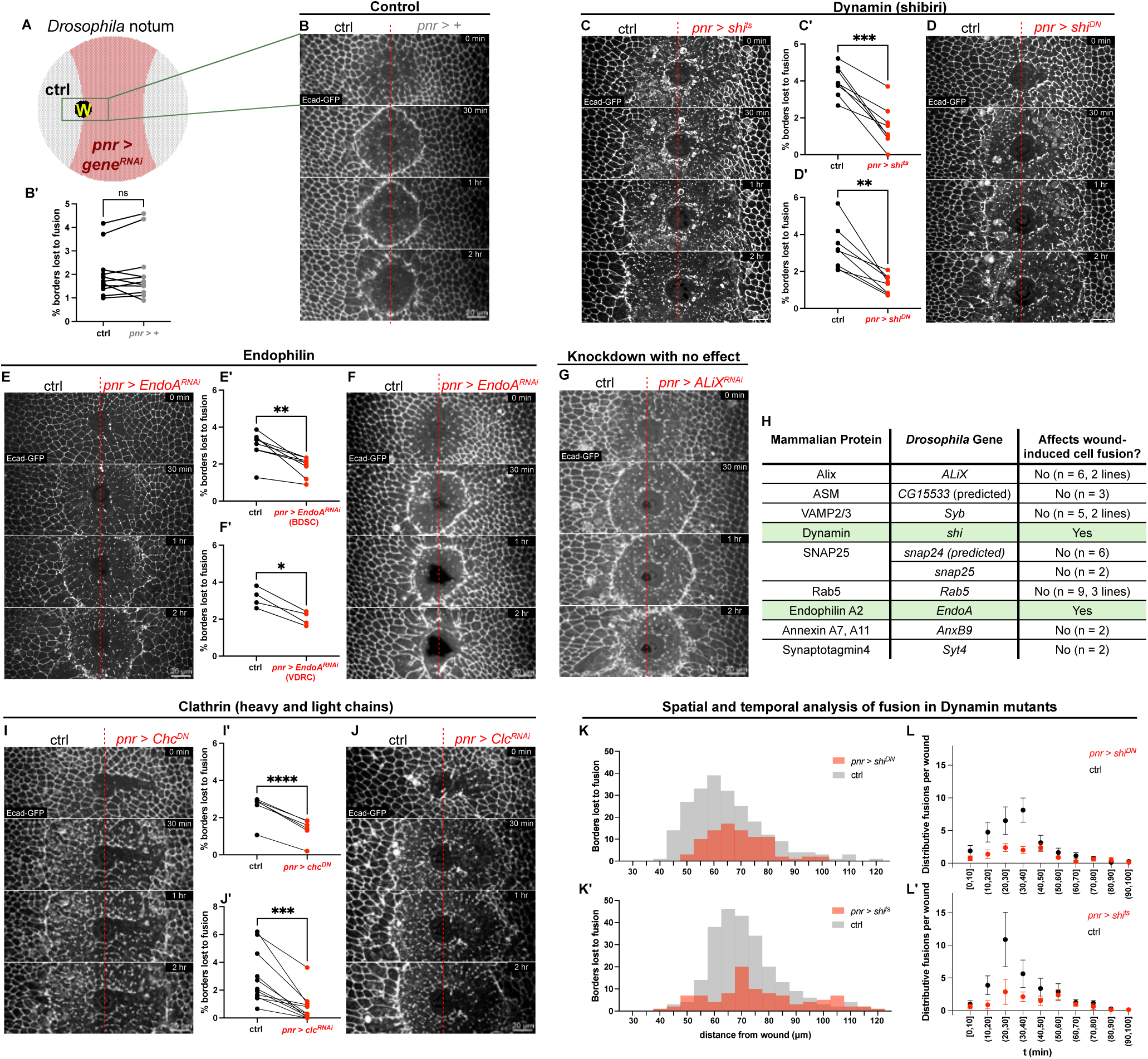
A candidate screen identifies protein regulators of wound-induced fusion. **A.** Schematic of the experimental system with internal control. Genes are knocked down in central *pnr* domain (pink). A laser pulse is targeted to the right or left edge of the *pnr* domain to make the wound (w). **B-B’.** In pupae without genetic manipulation, the frequency of fusion is comparable in the *pnr* and control domains: p = 0.57, paired T-test; n = 11 wounds in independent animals. **C-F.** A candidate genetic screen identified two endocytosis genes, *shi* and *EndoA*, whose inhibition decreased the frequency of wound-induced fusion: *shi^ts^*, n= 8, p= 0.0003 (paired T-test); *shi^DN^*,n= 8, p= 0.0015; *EndoA.RNAi* from BDSC, n = 8, p = 0.0022; and *EndoA.RNAi* from VDRC, n = 4, p = 0.012. **G.** An example negative result from the candidate screen: knocking down *ALiX* does not visibly change syncytia in the *pnr* domain. **H.** Table of candidate genes screened. **I-J. I**nhibition of a third endocytosis component, *clathrin*, also decreases the frequency of fusion: *chc^DN^*,n= 6, p< 0.0001; *clc.RNAi*,n= 11,p= 0.0004. **K-L.** The spatial and temporal profiles of wound-induced fusion with impaired or intact *shi* function in the *pnr* domain suggests that *shi* acts uniformly, regardless of distance from the wound or time after wounding. (n = 8, error bars are SEM).

Because wound-induced fusion is primed by plasma membrane damage, we targeted candidate genes involved in membrane repair and remodeling (Fig. 5H). Among the nine candidates tested by loss-of-function, most did not visibly alter the size of wound-induced syncytia (*e.g.* Fig. 5G), yet two substantially reduced syncytia size and fusion frequency: *shibire* (*shi*, the *Drosophila* gene encoding dynamin), and *EndophilinA* (*EndoA*). To confirm the specificity of *shi* loss-of-function, we tested two different conditions, a dominant-negative and a temperature-sensitive mutant: each decreased the frequency of fusion by ∼75% (Fig. 5 C,D). Likewise, knocking down *EndoA* with either of two different RNAi lines consistently reduced fusions by ∼50% (Fig. 5 E,F). We previously reported that cell fusion sometimes has a different morphological appearance: instead of losing adherens junctions first, a cell’s apical surface is reduced in area -- seeming to shrink -- as the cell fuses basally with a neighboring cell^1^. The loss of *shi* reduced the number of apically shrinking cells by ∼50-75%, similar to the reduction in fusion visualized by the loss of adherens junctions (Fig. S2). Given these results, we conclude that both dynamin and endophilin promote wound-induced fusion.

Identifying both dynamin (*shi*) and endophilin in our screen suggested an important role for endocytosis in wound-induced fusion, as dynamin’s proline rich domain binds to the SH3 domain of endophilin^62^ specifically during endocytosis. Dynamin is a GTPase that oligomerizes to form helices, and it has multiple functions: dynamin bundles actin filaments to promote actin-based protrusions^63^; and it forms helices around the neck of budding vesicles to facilitate their scission^64^, promoting endocytic budding. To test the endocytic role of *shi* in promoting fusion, we utilized a transgenic *shi* mutant specifically designed to be defective in endocytosis with a deletion of its pleckstrin homology (PH) domain. This deletion, denoted *shi.ΔPH,* has no effect on actin bundling, but it prevents oligomerization around lipid nanotubes^63^. We found that expression of *shi.ΔPH* decreased wound-induced fusion (Fig. S3), indicating that dynamin’s contribution to fusion relies on its endocytic function.

If endocytosis is required for wound-induced fusion, we reasoned that clathrin should also be required. Indeed, inhibiting clathrin with either of two different approaches, a dominant-negative transgene of *Clathrin heavy chain* (*Chc*) or RNAi against *Clathrin light chain (Clc)*, also decreased fusion frequency by ∼50% (Fig. 5 I,J). Hence, dynamin, endophilin and clathrin all promote wound-induced fusion through their coordinated roles in endocytosis.

### Dynamin promotes fusion, preferentially localizes to fusing membranes, and enhances wound healing

To characterize the spatial and temporal role for endocytosis in fusion, we further analyzed dynamin (*shi*) loss-of-function conditions by analyzing the late but visible fusion indicator, the loss of p120ctn. In unmanipulated controls (either *pnr* or control domain) the spatial profile of wound-induced fusion was broad, with a peak about 60-70 µm from the wound center and a long tail out to 120 µm (Fig. S4D). When *shi* was inhibited in the *pnr* domain (with *shi^ts^* or *shi^DN^*), fusion was reduced evenly, yielding a similar but flattened spatial profile compared to controls (Fig. 5K,K’). With respect to temporal requirements, in controls the loss of p120ctnRFP peaked between 20-30 min after wounding (Fig. S4E). When dynamin function was inhibited in the *pnr* domain (with *shi^ts^* or *shi^DN^*), fusion was reduced at all times, yielding a similar but flattened temporal profile (Fig. 5L,L’). Thus, dynamin promotes wound-induced fusion across the wound bed and throughout the duration of the fusion response after wounding.

One attractive model for how endocytosis promotes wound-induced fusion is that it specifically removes the plasma membranes lost between cells during fusion. This model predicts that dynamin would be specifically localized to those membranes, which we call fusion borders. To test this model, we tracked dynamin’s sub-cellular localization both before and during fusion. On expressing transgenic *shi-GFP* throughout the *pnr* domain, we observed that all cell borders were crowded with high levels of Shi-GFP even before wounding (Fig. S5), making it difficult to detect wound-induced dynamics. We next expressed *shi-GFP* in individual cells using Gal4-based FLP-out clones. In labeled cells, the high levels of Shi-GFP on cell borders before wounding still obscured post-wound dynamics (Fig. 6A). Fortunately, we were able to take advantage of the post-fusion transfer of Shi-GFP from a labeled cell to an unlabeled recipient cell, creating a recipient cell sparsely labeled with Shi-GFP. Among these sparsely labeled cells, a small fraction went on to fuse with secondary recipient cells, generating a fusion border (see schematic in Fig. 6B). We identified four such cases, and we focused our analysis on the localization of Shi-GFP with respect to the secondary fusion events in these sparsely labeled cells. In all four, Shi-GFP preferentially localized to the fusion border, which was subsequently involved in secondary fusion (as detected by loss of p120ctn). The other cell borders retained p120ctn and had consistently lower levels of Shi-GFP (Fig. 6C and Fig. S4A-C). Thus, after tissue injury, dynamin preferentially localizes to sites where cell fusion occurs.

**Figure 6:**
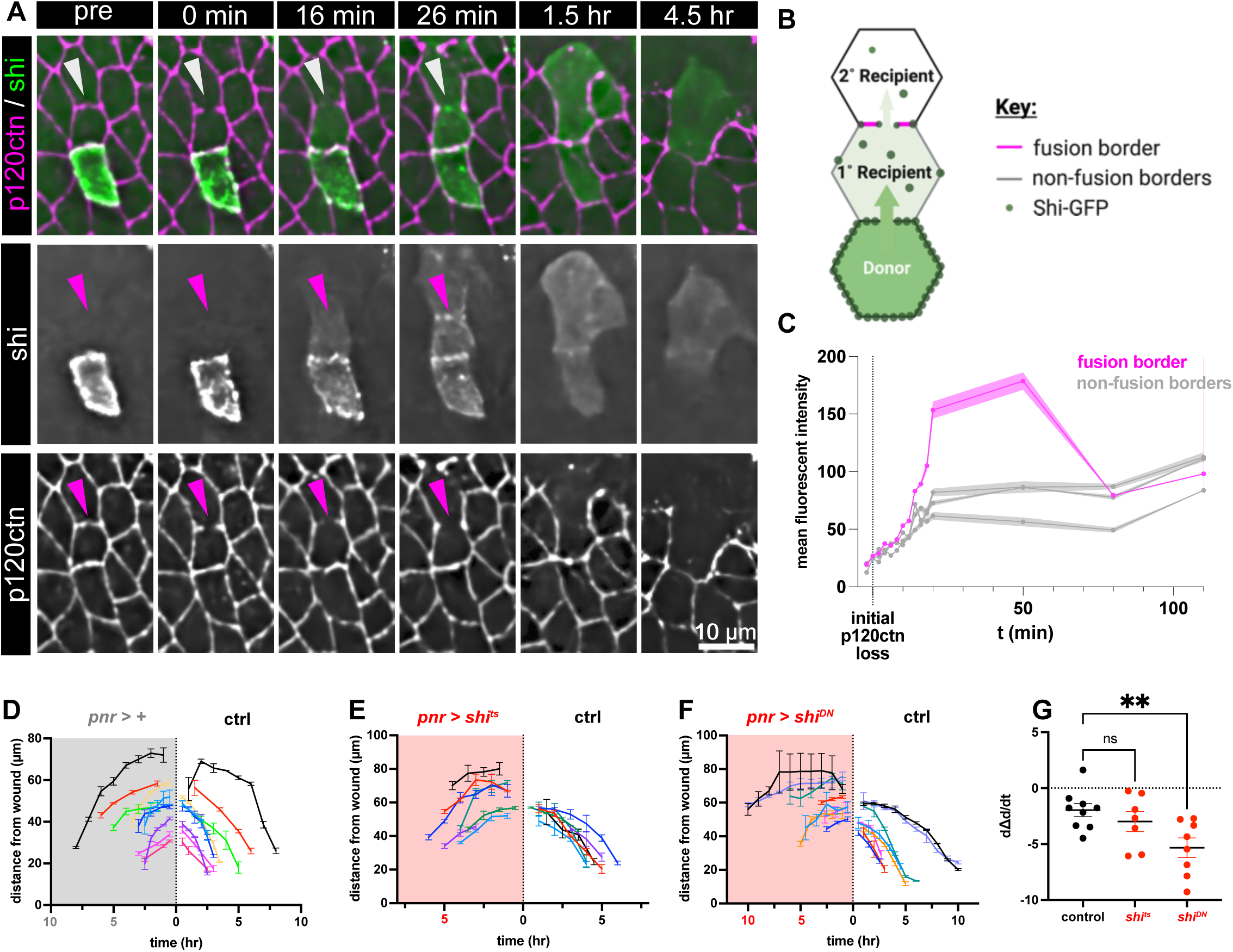
After wounding, dynamin localizes to cell borders where fusion occurs and speeds the rate of wound closure. **A.** Time-lapse images after a single wound showing that shi-GFP preferentially accumulates at sites of cell-cell fusion. This accumulation can be observed when a primary recipient fuses with a secondary recipient (arrowheads). **B.** Schematic of three cells: a donor, a primary recipient, and a secondary recipient. Cell borders of the primary recipient cell are labeled to indicate the fusion border, where Shi-GFP is recruited. **C.** Quantification of enrichment of shi-GFP along the primary recipient’s fusion border compared to non-fusing borders. Line thickness represents SEM. Similar preferential localization was observed in 4 of 4 samples (see Fig. S6). **D-F.** Wound closure rates assessed via time-dependent distance between the wound center and the leading edge in *pnr* and control domains. Without genetic manipulation, the leading edge advances slightly slower in the *pnr* domain (D). When dynamin is inhibited in the *pnr* domain by *shi^ts^*, this advancement becomes slightly more asymmetrical (E); and when dynamin is inhibited by *shi^DN^*, this advancement is much slower in the *pnr* domain where dynamin activity is reduced (F). **G.** Quantification of the speed of wound closure in control, *shi^DN^*-, and *shi^ts^*-expressing regions. While the speed of healing is comparable between control and *shi^ts^* regions (p = 0.57), it is significantly decreased in *shi^DN^* regions (p = 0.0093). *p* values calculated by one-way ANOVA with multiple comparison (mutants compared to the control); error bar denotes +/- SEM.

Since cell fusion has been shown to promote wound healing^8^, we quantified the speed of wound closure with and without dynamin inhibition. The distance from leading edge to the center of the wound was tracked in both the *pnr* (d_pnr_) and the control (d_ctrl_) domains (Fig. 6D-F). The difference in distances (Δd = d_ctrl_ - d_pnr_) over time was fit with linear regression, and the resulting slope (dΔd/dt) is the speed of wound closure (Fig. 6G). If the slope is negative, then the speed of wound closure is slower in the *pnr* domain: the more negative the slope, the slower wound healing is in the *pnr* domain relative to the control domain. <u>Even</u> without genetic manipulation the speed of wound closure is slightly slower in the *pnr* <u>domain</u> with mean <dΔd/dt> = -1.97 µm/hr (Fig. 6D,G). When dynamin function is inhibited by *shi^ts^*, the speed of wound closure is not significantly decreased compared to the controls (Fig. 6E,G, <dΔd/dt> = -3.00 µm/hr, *p* = 0.57, one-way ANOVA). However, dynamin inhibition by *shi^DN^* significantly impairs wound healing (Fig. 6F,G, <dΔd/dt> = -5.33 µm/hr, *p* = 0.0093, one-way ANOVA). The different effect on wound closure between *shi^ts^* and *shi^DN^*can be explained by their respective degree of inhibition of wound-induced fusion. Although the frequency of border-loss fusion is decreased to similar levels by the two mutants (Fig. 5 C-D), *shi^DN^* more severely inhibits shrinking fusion (Fig. S3 B-C) and has higher impact on overall wound-induced fusion (Fig. S3 B’-C’). Thus, cell-cell fusion speeds healing laser wounds in the pupal notum.

### Dynamin promotes fusion pore expansion

To further investigate the role of dynamin and endocytosis at sites of cell fusion, we carefully quantified the transfer of cytoplasmic GFP from donor to recipient cells in the hope of determining how dynamin influences the dynamics of fusion pores. Tracking the flow of cytoplasmic GFP between control donor-recipient pairs over five minutes, we observed that the time-dependent fluorescence levels were mirrored between donor and recipient: fluorescence lost in the donor was gained in the recipient (Fig. 7A-D). These data indicate that, even though cell pairs are connected by open channels at the earliest times, the loss of cytoplasmic GFP to the environment is minimal, confirming the earlier bystander analysis (Figs. 3B-D and S1).

**Figure 7:**
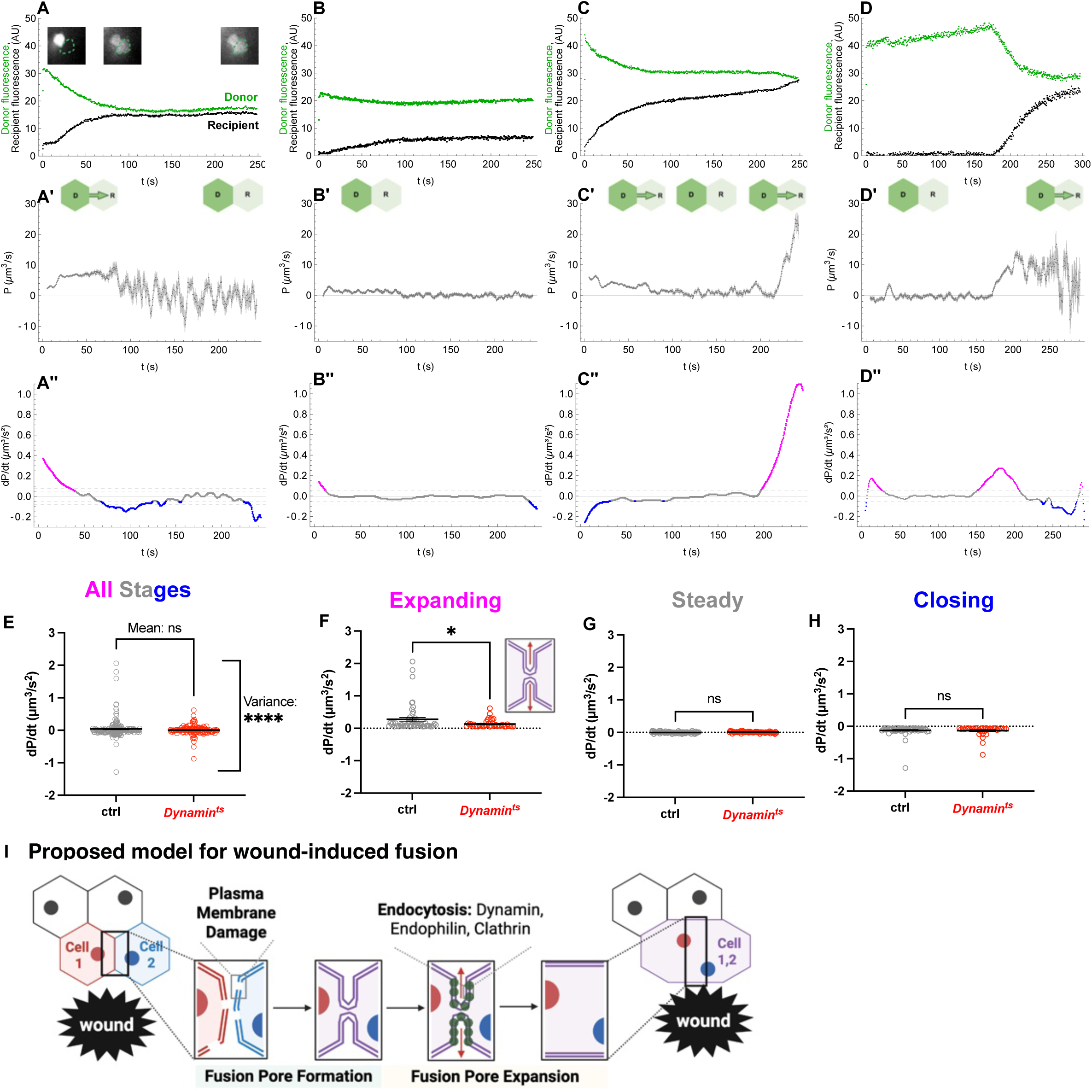
Dynamin is required in each fusion partner for effective fusion pore expansion. **A-D’’.** Analysis of the dynamics of cytoplasmic GFP transfer between four example pairs of donor and recipient cells. The fluorescence of each pair over time (A-D) was used to calculate both the dynamic permeance (A’-D’) and its rate of change (A"-D"). The four examples show GFP transfer that (A) reached equilibrium directly, (B) stopped before equilibration, (C) stopped and then re-initiated, and (D) did not initiate until > 150 s after wounding. **E-H**. Comparison of fusion dynamics between controls pairs and heterotypic pairs with *shi* function inhibited in only the donor cell. The rate of change in permeance (dP/dt) was used to categorize the state of the transfer channels at each time as expanding, steady or closing. When all three states were combined (E), the mean dP/dt was similar (p = 0.15, Welch’s T-test) but the variance was greater in control pairs (p < 0.0001). When the states were split, the mean dP/dt was reduced in heterotypic pairs specifically during the expanding state (F; p = 0.031, Mann-Whitney test) but not during steady (G; p = 0.091, Welch’s T-test) or closing states (H; p = 0.35, Mann-Whitney test), indicating a role for dynamin in promoting channel expansion. Bars denote SEM. **I.** Proposed model of wound-induced fusion: injury causes plasma membrane damage, which primes epithelial cells to form fusion pores, initiating the sharing of cytoplasm and plasma membrane (red/blue to purple); then, once pores are formed, endocytosis (green circles) promotes pore expansion to complete fusion.

Interestingly, the flow was not constant but rather underwent dynamic changes. In Fig. 7A, GFP transfer began about 10 s after wounding and continued uninterrupted until the cells equilibrated. In contrast, in Fig. 7B, the GFP levels started to converge, but then, about 95 s after wounding, they plateaued at different levels in the donor and recipient, suggesting that the channel had closed. A similar plateau is shown in Fig. 7C, but GFP transfer then resumes after a ∼2-min pause. These fluctuations in the rate of GFP transfer show that the channels between cells are not static, but that they undergo periods of expansion and contraction, and even closing and reopening. Interestingly, all open channels that we analyzed in control wounds, even those where GFP transfer stalled, eventually resumed GFP transfer and resulted in complete cell fusion (Fig. S7).

We then used this fluorescence data to calculate the time-dependent membrane permeance,

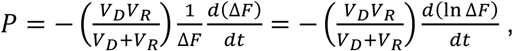

where *V_D_* and *V_R_* are the donor and receptor cell volumes, and Δ*F* is the fluorescence difference representing the concentration difference between donor and recipient. Given that multiple fusion pores are evident by TEM, the calculated membrane permeance represents the collective impact of all channels between two cells. Thus, permeance is a proxy for total fusion channel area^65^: higher permeance corresponds to a higher rate of GFP transfer for a given concentration difference and indicates that the channel area is larger; and reciprocally, smaller permeance indicates smaller channels. Examples of four control-pair permeance traces are shown in Fig. 7A’-D’, corresponding to the donor and recipient GFP levels in Fig. 7A-D. The dynamic changes in permeance are particularly notable in Fig. 7C,C’: as GFP transfer arrests from about 95 to 215 s and then resumes, the permeance correspondingly drops from ∼5 µm^3^/s to near zero over the first 95 s, bounces around near zero for the next 120 s, and then rapidly ramps up to over 10 µm^3^/s just before the cells equilibrate.

To determine how dynamin function impacts fusion pore dynamics, we compared the time rate of change of permeance, *dP/dt*, in control pairs and in cases where dynamin function was reduced via *shi^ts^*. Since *Gal4* was used to drive both cytoplasmic GFP labeling and expression of *shi^ts^*, we were not able to reduce dynamin function in both donor and unlabeled recipient; our comparison is thus limited to homotypic control-control and heterotypic *shi^ts^*-control fusion pairs. As shown in Fig. 7A"-D", GFP transfer between a specific pair of cells can have periods of both increasing and decreasing permeance: positive values of *dP/dt* (shown in magenta) indicate periods when channels or pores are expanding, whereas negative values of *dP/dt* (shown in blue) indicate when they are contracting. For control-control pairs, the average rate of permeance change was biased towards pore expansion (<*dP/dt* > = +0.04 µm^3^/s^2^) but with substantial variance (s.d. = 0.28 µm^3^/s^2^). A few intervals even had a rate over 1 µm^3^/s^2^ (Fig. 7E). The average rate of permeance change for heterotypic *shi^ts^-*control pairs was not substantially different (<*dP/dt* > = +0.007 µm^3^/s^2^; *p* = 0.19, Welch’s t-test), but the variance was much smaller (s.d. = 0.14 µm^3^/s^2^; *p* < 0.0001, Welch’s t-test). As Fig. 7E shows, the primary difference was that heterotypic *shi^ts^* pairs were missing intervals with very large *dP/dt* values (> 1 µm^3^/s^2^), *i.e.*, they were missing periods of rapid pore expansion.

To quantify this effect, we separately compared the three interval types – expanding, steady, and closing. Loss of dynamin function did not affect the time spent in each of the three stages (Fig. S8A-C), but it did decrease the speed of fusion pore expansion (*p =* 0.045, Mann-Whitney test; Fig. 7F). No significant differences were observed in the steady state (*p =* 0.063, Welch’s t-test) or closing intervals (*p =* 0.42, Mann-Whitney test). Thus, inhibiting dynamin function, even in just one member of a heterotypic donor-recipient pair, reduced the mean rate of fusion pore expansion, most notably apparent at the upper end of the expansion-rate distribution. To rule out the possibility that this reduction was a result of biased sampling, we confirmed that our samples of control and heterotypic *shi^ts^*-control pairs were at comparable distances from the wound (Fig. S8D, D’) and had similar levels of tissue damage (Fig. S8E, E’). Dynamin thus promotes cell fusion by increasing the rates at which fusion pores expand.

## Discussion

### A four-step model of wound-induced fusion

Integrating our data, we propose the following four-step model of wound-induced cell-cell fusion: (1) the wound damages the plasma membranes of some cells that nonetheless survive, priming those cells to fuse; (2) after provisional membrane repair, unstable fusion pores are established at sites of membrane damage anywhere from minutes to hours after damage; and (3) expansion of these fusion pores is promoted by endocytic removal of excess membrane at the fusion border until (4) adherens junctions are removed and two distinctly visible cells merge into one (Fig. 7I).

The first step, i.e., the priming of fusion by wound-induced plasma membrane damage, is supported by the observation that every cell border removed by fusion had previously been rendered transiently permeable to extracellular Ca^2+^, as early as 30 ms after wounding. Based on our prior finding that the early phase of Ca^2+^ entry correlates with dye entry^53^, we attribute these very early Ca^2+^ influxes to membrane damage rather than the opening of any specific ion channel. The early presence of plasma membrane damage is further supported by our observations that transfer of cytoplasmic GFP from donor to recipient cells begins as early as 500 ms after wounding. Finally, we establish a causative and not just correlative link by showing that cell fusions no longer occur when wounds are made without collateral plasma membrane damage.

The second step, i.e., the conversion of membrane damage sites to membrane-lined fusion pores, is supported by the observation that the sharing of cytoplasmic GFP within seconds-to-minutes of wounding is a reliable indicator that a cell pair will eventually fuse; and yet, free lipid diffusion between cell pairs, indicated by the transfer of transmembrane mCD8-GFP, was evident only after a delay of several minutes and could take much longer. Given the length of this delay, it is likely that formation of the fusion pore was preceded by provisional membrane damage repair^60^. In fact, for several cell pairs, the time course of cytoplasmic GFP sharing follows a start/pause/restart pattern that matches expectations for damage/provisional repair/fusion pore formation. Finally, we used TEM of fixed samples to directly image multiple fusion pores between adjacent cells.

The third step, i.e., the expansion of fusion pores driven by endocytic removal of adjacent membrane, is supported by our genetic screen, dynamin localization, and our quantitative analysis of the dynamics of fusion pore permeance. The screen directly identified two endocytosis genes that promoted wound-induced fusion – dynamin and endophilin A – and based on those hits, we further tested and confirmed a fusion-promoting role for the endocytosis genes encoding clathrin heavy and light chains. To further characterize the role of endocytosis in fusion, we analyzed the localization of dynamin with respect to fusion, finding that dynamin is preferentially targeted to cell interfaces where complete fusion will later occur, consistent with a role for endocytosis in expanding fusion pores. Analyzing fusion pore permeance, we determined that fusion pores are highly dynamic, fluctuating in diameter over time, but with a bias toward expansion; inhibiting dynamin’s function reduces the rates of pore expansion.

Previous studies reported that autophagy is also required for wound-induced cell fusion^33^, and we speculate that autophagy promotes recycling of endocytosed membrane necessary to complete fusion. Finally, the fourth and the last visible step in cell fusion is the removal of adherens junctions between the fusing cells, observed directly and used to denote the time at which two cells merge into one morphological syncytium.

### Plasma membrane damage may act as a fusogen

Our fusion mechanism may seem to lack a key player – a protein fusogen that overcomes the energy barrier to bring opposing membranes into proximity. Unlike developmentally fusing cells that are genetically pre-programmed to fuse by expressing a fusogen, we expect that canonical fusogens are not readily available to non-fusogenic epithelial cells responding to an abrupt and unforeseen wound. What can a wounded cell use to promote fusion pore formation? We propose that damaged plasma membranes act directly as fusogens. Under the definition proposed by Bruckman et al., fusogens must meet three criteria: (1) be observed on fusion borders at the time of fusion; (2) be required for fusion; and (3) be able to induce ectopic fusion among non-fusogenic cells^36^. Plasma membrane damage meets all these criteria.

We can envision many potential mechanisms for plasma membrane damage to induce ectopic fusion. The most obvious is that damage opens the plasma membrane, which could then reseal with membrane from another cell; however, this mechanism is not consistent with either the delayed initiation of mCD8 sharing nor the large fraction of late cytoplasmic-transfer events that occur minutes-to-hours after wounding, because it requires cells to remain open to the environment for minutes to hours, and cells are expected to reseal on a much faster timescale^60^. One possibility is a variant of this model that accommodates the longer timescales by having damaged plasma membranes form a hemifusion-stalk, where the outer leaflets of two plasma membranes fuse while the inner leaflets remain separate. A rapidly formed hemifusion intermediate could later be resolved either by fusion of the inner leaflets into a fusion pore, or by fusion of the outer leaflet into separate cells. A second possibility focuses on provisionally repaired plasma membrane, where membrane-bound organelles relocate to the site of damage and form a temporary patch to close the breach^66^. Although this patch would be slowly remodeled to restore the plasma membrane to its undamaged state, we propose that the patched membrane could be fusogenic: if two patches come in contact into each other, they could intermingle and repair as one cell rather than two. In this model, the variable time observed for the initiation of sharing could be explained by stochastic sliding of membrane patches, which would "bump into" each other over a wide range of time scales. A third possibility focuses on the integral membrane protein complexes that connect cells, such as adherens junctions, gap junctions, and tight (septate) junctions. Wound-induced damage could denature and aggregate junctional proteins in the extracellular environment, creating a stable linkage between cells that would need to be removed by endocytosis. However, the uptake of aggregated junctional proteins risks simultaneously endocytosing some of the facing plasma membrane, leading to fusion. We note that this model requires endocytosis for both fusion pore formation and expansion. Although the mechanism remains unclear, our data indicate that plasma membrane damage is a non-protein fusogen that primes cells for fusion after tissue injury.

### Plasma membrane damage is common in wounds

Although the studies here exclusively investigate laser wounds, plasma membrane damage occurs in many types of naturally-occurring abrasion and puncture wounds^67^. It also occurs following mechanical rupture of cells within the body such as when cancer cells squeeze through narrow openings in the surrounding matrix during metastasis^68^. Further, injuries that cause plasma membrane damage also frequently yield cell-cell fusion – e.g., in response to laser^1,33^, puncture and pinch wounds in *Drosophila*^8,69,70^, in response to muscle injuries in vertebrates where damaged muscle cells fuse with satellite cells^2^, and after membrane-damaging electric pulses have been applied to *Dictyostelium discoideum*^71^. Despite these examples, it is difficult to definitively detect cell fusion without live cell imaging, and its occurrence during tissue repair can be easily overlooked. In fixed samples, the presence of multinucleated syncytia may represent either cell fusion or a failure of cytokinesis; and the absence of syncytia does not exclude cell fusion, because wound-induced syncytia may have formed but then migrated away or been eliminated.

Despite the difficulty in detecting cell fusion after injury, previous literature^8^ and our data show that it is important for wound healing. Cell fusion can facilitate tissue repair in multiple ways^72^. For example, homotypic cell fusion allows the sharing of proteins, organelles and chromosomes among damaged cells to form a healthier reservoir of cellular contents, rescuing cells that may otherwise die. There are reports of cell fusion successfully introducing wild-type alleles of disease genes into mutant mice, including *Fah* in *Fah^-/-^* mice^19–21^ and dystrophin in *mdx* mice^12^. In addition, heterotypic post-injury fusion between stem cells and resident somatic cells promotes the restoration of biomass, either by stem cells adopting the profile of somatic cells^73–76^ or by the reprogramming of somatic cells to re-enter the cell cycle^77–85^. Such cell-cycle re-entry is also the mechanism by which heterotypic cardiomyocyte-endothelial cell fusions help repair damage to cardiac tissue^86^. Thus, cell-cell fusion in the context of tissue injury represents a fascinating problem in cell biology and offers potential clinical therapies.

## Acknowledgements

We thank Rachel C. Hart and Maria Vingradova for assistance with the transmission electron microscopy sectioning, and J. White, K. LaFever, A. Stricker, and members of the Page-McCaw lab for general help and support. We thank Dr. E. Chen (University of Texas Southwestern Medical Center, Dallas, TX) for suggesting that we test clathrin and for providing the *shi* stocks; M.I. Prislusky from Stephanie Seveau’s lab at Ohio State for suggesting a plasma membrane repair genes for us to screen; and the Bloomington Drosophila Stock Center (Bloomington, IN) and Vienna Drosophila Stock Center for *Drosophila* stocks. Transmission electron microscopy was performed in part through the use of the Vanderbilt Cell Imaging Shared Resource (supported by NIH grants CA68485, DK20593, DK58404, DK59637, EY08126, S10OD034315, R24OD037694, and S10MH137068). This work was funded by the National Institute of Health (R01GM130130 to A.P.-M. and M.S.H.) and the American Heart Association (25PRE1374646 to J. H.).

## Materials and Methods

### Live imaging and pupal dissection

Pupae were dissected and mounted for imaging using previously published methods^57^. Live imaging was conducted on a Nikon Ti2 Eclipse with X-Light V2 spinning disk (Nikon, Tokyo, Japan) as described previously^1^. Its “Fast time-lapse imaging” function was used to obtain movies with intervals of 30-ms (for fast calcium signal) or 500-ms (for immediate GFP sharing). A microscope heating stage and an objective heating collar were used to heat some of the samples. After acquisition, movies were processed with NIS-Elements AR (version 5.30.05). All wounding and most imaging were done at 40x, aside from mCD8 clones imaged at 40x with 1.5x zoom. The pixel-to-µm scale was 0.28 µm/pixel.

### Laser ablation

Pupae were wounded using previously described protocols for single-pulsed ablation^57^ and scanning ablation^56^ using a nanosecond UV laser (3rd harmonic of a Q-switched Nd:YAG) focused at the optical plane containing fluorescently-labelled p120ctn or E-cadherin. For scanning ablation, the same area was scanned/wounded three times to thoroughly eliminate live cells. Laser pulse energy was adjusted as needed to maintain comparable levels of damage among different samples. The target damage levels were removal of E-cadherin signal throughout a circular region of radius 20-30 µm and/or loss of nuclear-confined nuc-mCherry signal, indicative of nuclear membrane damage, within one min of ablation out to a radius of ∼70 µm. The laser pulse energy used for scanning ablation was typically about 0.27 of that used for single-pulse wounds.

### Image analysis protocols

To generate profile plots of 30-ms GCaMP6m and p120ctn signals (Fig. 1E), the “Straight” tool in FIJI was used to draw a 10-pixel wide line out radially from the center of the wound. Intensity along the line was extracted using the “Plot Profile” function and plotted in Prism. To determine if a specific fusion cell border co-localized with sites of plasma membrane damage (Fig. 1F-H), the manually annotated fusion cell border was overlain on the 30-ms GCaMP6m signal, and co-localization was determined visually.

To determine whether the 30-ms calcium/GCaMP6m signals were affected by genetic manipulations in the *pnr* domain, we surveyed three individuals who were blinded to sample identity. Each participant was presented with a random shuffling of the 30-ms GCaMP6m images from three conditions (*pnr > +, pnr > shi.ts, pnr > EndoRNAi*). Given no information about genotype or domain identity (*pnr* could be on the left or right), each participant was asked to identify whether the calcium signal had spread further on one side of the wound or the other.

To determine whether the region of cell lysis was affected by genetic manipulations in the *pnr* domain, we quantified the area of lysis in the *pnr* and control sides at 30 min and 1 hr after wounding. Quantification used FIJI’s “Freehand Tool” to manually draw ROIs that passed through the targeted wound location and traced the accumulation of E-cadherin signal along the leading edge on each side of the wound. The area of each ROI was generated using FIJI’s “Measure” function.

To track the intensity of shi.GFP on fusion and non-fusion cell borders (Fig. 6C, Supplement 4A-C), we collected two-color z-stacks with p120ctn.RFP and shi.GFP. Since cells in the notum are highly non-prismatic, analysis was limited to the single z-slices where p120ctn was manually determined to be in focus. The isolated p120ctn.RFP channel was used to draw an ROI for each cell border to be analyzed (drawn with FIJI’s “Straight” tool at a width of 3 pixels). Those ROIs were then applied to the shi.GFP channel to extract the mean and standard deviation of the shi.GFP intensity for each border over time (using FIJI’s “Measure” function). Based on the number of pixels within the ROI, standard deviation was converted to standard error of the mean (SEM). Note that the separation and isolation of channels was applied to avoid having the shi.GFP signal bias the drawing of ROIs.

### Fusion analysis

A cell border was counted as a location of wound-induced fusion if it met three criteria: (1) adherens junctions along the cell border were lost; (2) the lost adherens junctions were not restored; and (3) the loss of adherens junctions was accompanied by a morphological change of the neighboring cells, for example, the moving apart of tri-cellular junctions previously connected by the lost cell border. Since border-breakdown fusions are observed most frequently within 1 hr after wounding^1^, we counted all such fusions occurring within 1 hr 40 min of wounding.

### Analysis of fusion frequency

In wild-type samples, the frequency of fusion was calculated as the number of fusion events observed within a given area divided by the estimated number of cell borders in that area. To encompass all wound-induced fusions, the observed area was an annulus with its inner radius matching the cell lysis zone (observed 6 min after wounding) and its outer radius being 380 pixels (106 µm). The number of cell borders in that annular region was estimated by area-based scaling of the border count within a randomly selected 70-x-70-pixel (19.6-x-19.6-µm) region in the pre-wound image.

When comparing the fusion frequency in *pnr* versus control domains, we modified the observation area to a landscape-oriented rectangle centered on the wound with its height defined by the inner radius of the zone of nuclear membrane damage (approximately the same as the zone of lysis). Replacing the annular region with an extended rectangle excludes possible heterotypic fusions along the *pnr*-control border just above and below the wound. The total number of cell borders was estimated for each side based on a similar sampling method as above. To ensure random region selection and eliminate potential human bias, we made sure that the screen’s least significant hit (*EndoA RNAi*) had its sample region coordinates selected using pseudo-random numbers generated in a terminal command line using the python function random.randint (start, end).

The time of each border breakdown was recorded as the frame when one could first observe discontinuity in that border’s adherens junctions. The distance from the wound center was calculated based on the location where the laser was targeted (center of the image) and the position of the fusion cell border in the pre-wound or 0-min post-wound image, as marked using FIJI’s “Multi-point” tool.

Since fusions can also occur via apical shrinkage rather than border breakdown^1^, we also quantified the number of shrinking cell fusions in *pnr* versus control domains. Shrinking cell fusions were defined as diploid cells that decreased in surface area over time and fully disappeared within a 5-hr movie. As with fusion analysis, cells near the *pnr*-control border were excluded. Shrinking cells were counted manually and their spatial locations marked using FIJI’s “Multi-point” tool.

### Determining the speed of wound closure

The distance between the leading edge and the wound center over time was used to calculate the speed of wound closure. The distance was calculated using the coordinates of the wound center and chosen points on the leading edge. To choose positions on the leading edge so that the choice is unbiased and consistent over time, four reference lines, all crossing the center of the wound, were drawn using the red channel with only the nuc-mCherry signal: line 1 along the edge of the *pnr* domain labeled by nuc-mCherry, line 2 perpendicular to line 1, and line 3 and 4 are 30° from line 2. These references lines were then overlaid on the green channel with the E-cadherin signal. The intersection of E-cadherin marked leading edge and reference lines 2-4 were used as the position of the leading edge. That is, at each time point, a total of 6 positions of the leading edge were recorded, among which 3 will be in the *pnr* domain and the other 3 in the ctrl domain. The average distance between the leading edge and the center of the wound were calculated at each time point for both the *pnr* (d_pnr_) and the control (d_ctrl_) domain.

The difference between control and *pnr* distance (Δd = d_ctrl_ - d_pnr_) was calculated over time, and the resulting Δd over time (t) plots were fit to straight lines using Nonlinear regression function of Prism. The slope of the line (dΔd/dt) was the speed of wound closure, and the slopes were compared among control, shi.ts and shi.DN samples using one-way ANOVA with multiple comparisons (comparing the mean of each column with the mean of the control column) to generate the *p* value.

### Determining timescales for cytoplasmic GFP sharing

To determine the earliest time of cytoplasmic GFP sharing, donor-recipient pairs were selected for analysis based on post-wounding movies covering a duration of 5 min at a 500-ms frame interval. GFP intensity in the recipient cell and a neighboring bystander cell were quantified using FIJI’s “Measure stack” function. A recipient cell was considered to have acquired GFP within 500 ms of wounding if it met three criteria: (1) its intensity was already higher than that of a bystander cell within 500 ms; (2) its intensity remained higher than the bystander; and (3) its intensity continued on a general upward trend.

To determine the temporal distribution of later cytoplasmic GFP sharing events, we analyzed post-wound movies with either 2-min or 30-min frame intervals. To avoid parallax issues, analysis was limited to single z-slices at the level of the adherens junctions. The time of sharing initiation was recorded as the frame when an unlabeled cell first visibly acquired GFP. Note that we counted all visible recipient cells, even if a unique donor was not obvious, which means that the count includes not only primary recipients, but also secondary and potentially tertiary recipient cells – i.e., those that acquired cytoplasmic GFP from earlier recipients. The temporal distribution of cytoplasmic GFP sharing events was characterized by compiling the time-dependent cumulative count of GFP sharing initiations (in the 2-min interval movies) and using a python script to fit that data to an exponential approach to saturation. The time constant from this fit is the mean time for the initiation of cytoplasmic GFP transfer after injury.

### Determining timescales for membrane-tethered GFP sharing

To profile the temporal distribution of mCD8-GFP sharing, we acquired two types of movies within the first 5 min after wounding – either fast time lapse with 500-ms frame intervals to capture any immediate sharing or continuous z-stacks to capture any basolateral sharing. After this 5-min period, post-wound z-stacks were acquired at 2-min intervals out to 30 min. Analysis was performed as above for cytoplasmic GFP sharing but limited to primary recipient cells and using z-stacks with adjusted image gain to pick up mCD8-GFP’s fainter signal. When GFP was visible in a recipient cell for two consecutive frames, whether apically or basolaterally, the time of the first frame was recorded as the time of mCD8-GFP sharing initiation.

### Cytoplasmic GFP permeance analysis

This analysis was derived from Nolan et al^65^ and applied to donor-recipient pairs observed for 5 min after wounding at 500-ms frame intervals. The spatially averaged fluorescence intensity was measured for each donor and recipient cell using FIJI’s “Rectangle” tool to draw ROIs that enclosed a representative area within each cell and stayed within that cell as it moved. Once ROIs were drawn and saved, FIJI’s “Measure Stack” function was used to collect the average fluorescence intensity and its spatial standard deviation within each ROI at each time point. The time-dependent fluorescence background was acquired using a similar process applied to a neighboring unlabeled region and subtracted from the donor and recipient fluorescence.

The time-dependent permeance, *P*, for each donor-recipient pair was then calculated via

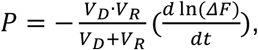

where *ΔF* is the time-dependent difference in GFP fluorescence intensity between donor and recipient, assumed to be proportional to the concentration difference, and where *V_D_* and *V_R_* are the estimated volume of each donor and recipient cell. The relative volumes were estimated using the apical surface area outlined by p120ctn. The numerical derivative 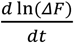 was calculated at each time point using a second-order polynomial fit to a sliding window of 21 centered datapoints. Thus, for a 500-ms frame interval, the window used to estimate the derivative spans a 10-s interval.

To calculate the rate of change in permeance, 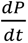, the numerical derivative was estimated at each time point using a second-order polynomial fit to a dynamic sliding window of 121 points. The wider window was needed to suppress noise in the calculation of what is a second derivative of the raw data. The window was dynamic in the sense that it was asymmetrically truncated for the first and last 60 permeance values where a full centered window could not be taken. For example, to estimate 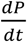 at the earliest time point, only the first 61 permeance values were included in the window; the slope of the fit was then calculated at the earliest time point rather than the center of the window. The resulting list of time-dependent 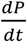 values was separated into intervals of fusion pore expansion, closing and steady state using a hysteresis filter with two upper thresholds (0.05 and 0.08 µm^3^/sec^2^) and two lower thresholds (-0.05 and -0.08 µm^3^/sec^2^). The length of each interval and the mean of 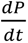 within each interval were calculated and compared between control clones and *shi.ts* clones.

### Transmission Electron Microscopy (TEM)

To prepare wounded samples for sectioning and TEM, pupae expressing shg.GFP were aged to 15-18 hr APF at 25°C and mounted following previously published methods^87^. To increase the probability of sectioning into a wounded region in later steps, four or five laser ablations were performed along the midline of each sample’s dorsal notum, with each wound ∼100-150 µm apart.

Samples were unmounted ∼30-50 min after wounding, and each pupa was then completely removed from its pupal case. To improve fixation without disturbing the area of interest, a scalpel was used to cut the ventral surface of each pupa through to the middle of the abdomen. Samples were fixed by submersion in 4% paraformaldehyde, 3% glutaraldehyde in 0.1 M cacodylate for ∼24 hrs. After primary fixation, the samples were cryoprotected by gradually transitioning them to 30% glycerol in 50 mM HEPES over 24 hours. Samples were vitrified using a Leica EM-ICE high-pressure freezer using two B hats for maximum depth. After vitrification, samples were freeze substituted in 1% uranyl acetate in absolute methanol at -90° C for 48 hours and then transitioned to -20° C over the next 24 hours. Samples were washed in methanol and transitioned to rehydration using the following scheme: 70% methanol at -20° C for 1 hour; 50% methanol at -20° C for 30 min; 30% methanol for 30 min at 4° C; 10% methanol at 4° C for 30 min; 0.05 cacodylate at 4° C for 10 min; and finally to 0.1 M cacodylate buffer at 4° C. Samples were then subjected to an OTO process: incubation with 1% OsO_4_, 0.75% potassium ferrocyanide in 0.1 M cacodylate for 1 hour; followed by 1% thiocarbohydrazide for 20 minutes; and then 2% OsO_4_ for 40 minutes Samples were then *en bloc* stained with 1% uranyl acetate for 1 hour and then dehydrated through a graded ethanol series. Samples were finally infiltrated with Epon-812 using propylene oxide as the transition solvent; and the infiltrated resin was polymerized at 60° C for 48 hours. Sections were cut to a nominal thickness of 70 nm on a Leica Enuity ultramicrotome and post-stained with 2% uranyl acetate and lead citrate. All samples were imaged on a JEOL 2100+ transmission electron microscope (JEOL, Peabody, MA) operating at 200 keV using a NanoSprint15 MK-II CMOS camera (AMT, Woburn, MA). Tilesets were acquired using SerialEM and reconstructed using the etomo/IMOD software suite.

### Fly husbandry

Most fly lines were obtained from the Bloomington Drosophila Stock Center: *w^1118^, UAS-ArcLight* (BDSC 51056), *UAS-ALiX.RNAi* (BDSC 50904, BDSC 33417), *UAS-CG15533.RNAi* (BDSC 36761), *UAS-Syb.RNAi* (BDSC 44014, BDSC 39067), *UAS-snap24.RNAi* (BDSC 28719), *UAS-snap25.RNAi* (BDSC 27306), *UAS-AnxB9.RNAi* (BDSC 38523), *UAS-Syt4.RNAi* (BDSC 26730), *UAS-shi.DN* (BDSC 5822), *UAS-shi.ts* (BDSC 44222), *UAS-EndoA.RNAi* (BDSC 27679), *UAS-chc.DN* (BDSC 26874), *UAS-clc.RNAi* (BDSC 27496), *UAS-mCD8.GFP* (BDSC 32184), *ubi-shg.GFP* (DGRC 109007). One line was obtained from the Vienna Drosophila Stock Center: *UAS-EndoA.RNAi* (VDRC 24617). Several fly lines were generated in our lab: *p120ctn.RFP, ActinGCaMP6m/CyO. shg.GFP; pnr-Gal4, tub-Gal80^ts^, UAS-nucmCherry/SM6-TM6B. p120ctn, AyGal4/CyO. hsflp, UAS-GFP/TM3*.

To inactivate *UAS-shi.ts*, the microscope heating stage and the collar wrapped around its 40x objective were both heated to 31°C. When *UAS-shi.ts* was overexpressed within the *pnr* domain, mounted slides were pre-heated on the microscope for 5-10 min before imaging. When UAS-shi.ts was overexpressed in clones, slides were pre-heated on the microscope for 10-30 min before imaging.

To induce FLP-out clones, 3^rd^ instar larvae were picked from crosses grown at 25° C. The larvae were then transferred to an empty vial so that heat shock could be done in a 37° C water bath. Finally, the heat shocked larvae were moved to a vial with food and allowed to grow at 25° C until they evert and reach the developmental stage for analysis.

*UAS-shi.GFP* used in Zhang et al.^63^ and shi^2^; UAS-shi.ΔPH.GFP/CyO were generously gifted by Dr. Elizabeth Chen and her lab.

Because *shi* is on the X chromosome, virgin *shi^2^; UAS-shi. ΔPH.GFP/CyO* was crossed to male *shgVenus; pnrGal4, tubGal80ts, UAS-nucmCherry / SM6-TM6B*, and grew at 18° C. Non-tubby 3^rd^ instar larvae were switched to 29° C until they evert and reach the developmental stage for analysis. Pupae with GFP signal were used for wounding and analysis.

To overexpress *UAS-RNAi* lines by inactivating Gal80^ts^, the various RNAi lines were grown at 18° C before being moved and kept at 29° C for various hours as listed: *UAS-shi.DN* for 7.5-13 hr; *UAS-shi.ts* for 8.5-13 hr; *UAS-EndoRNAi* (BDSC 27679) for 1-5 days; *UAS-EndoRNAi* (VDRC 24617) for 3-5 days; shi^2^; UAS-shi.ΔPH.GFP/CyO for 17-32 hr; *UAS-chc.DN* for 3-5 days; *UAS-clc.RNAi* for 3-5 days; *UAS-ALiX.RNAi* for 20-36 hr.

---

**Figure S1:**
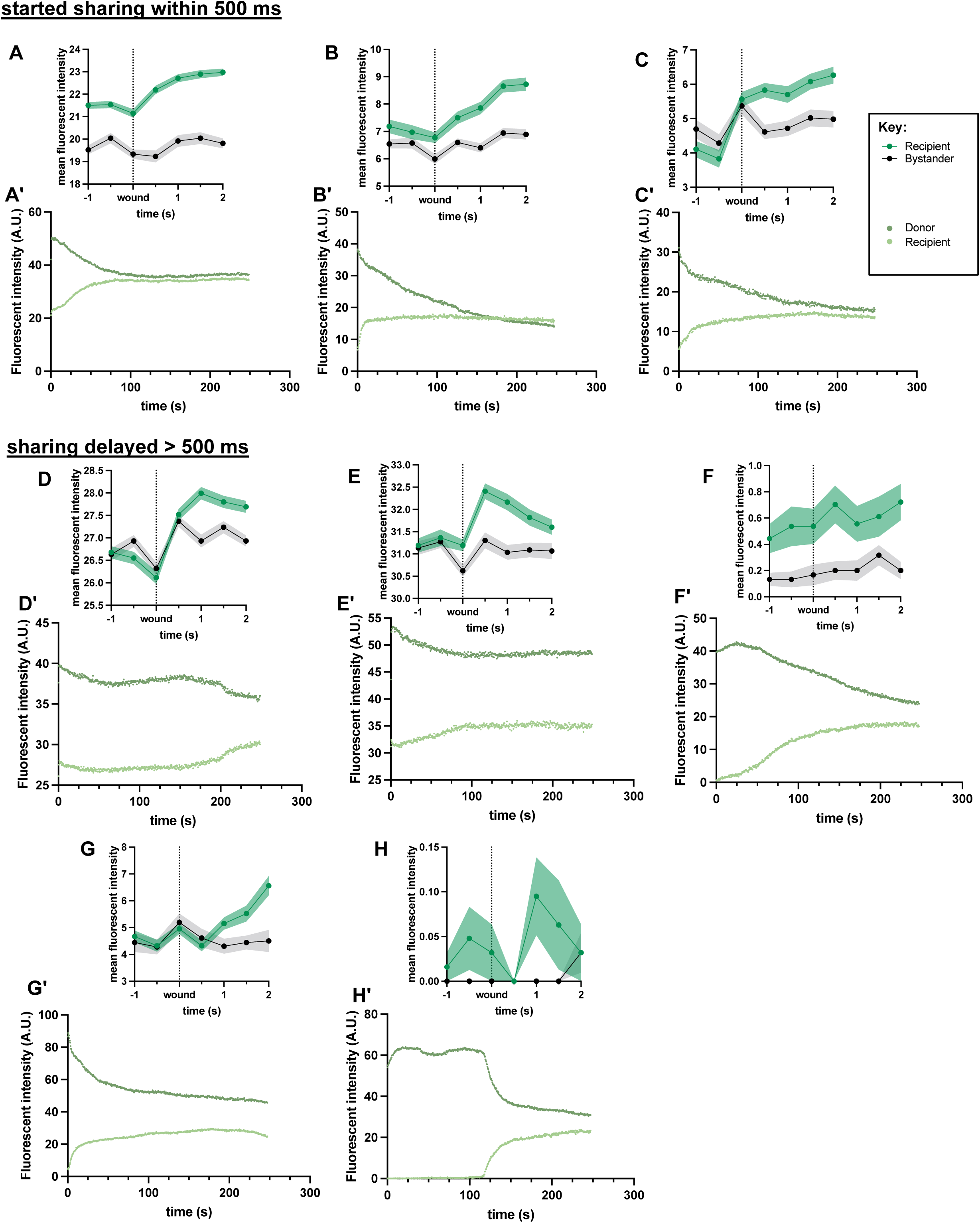
Additional analyses of fusion initiation time. Ten fusion initiation events were analyzed; two shown in Fig. 3, the remaining eight shown here. A-C. Three fusion events with GFP transfer initiated within 500 ms in addition to the one shown in Fig. 3C, used to quantify Fig. 3E. D-H. Five fusion events with GFP transfer initiated after 500 ms in addition to the one shown in Fig. 3D, used to quantify Fig. 3E. A’-H’. GFP levels over five minutes, in both donor and recipient cells for each fusion event, corresponding to panels A-H.

**Figure S2:**
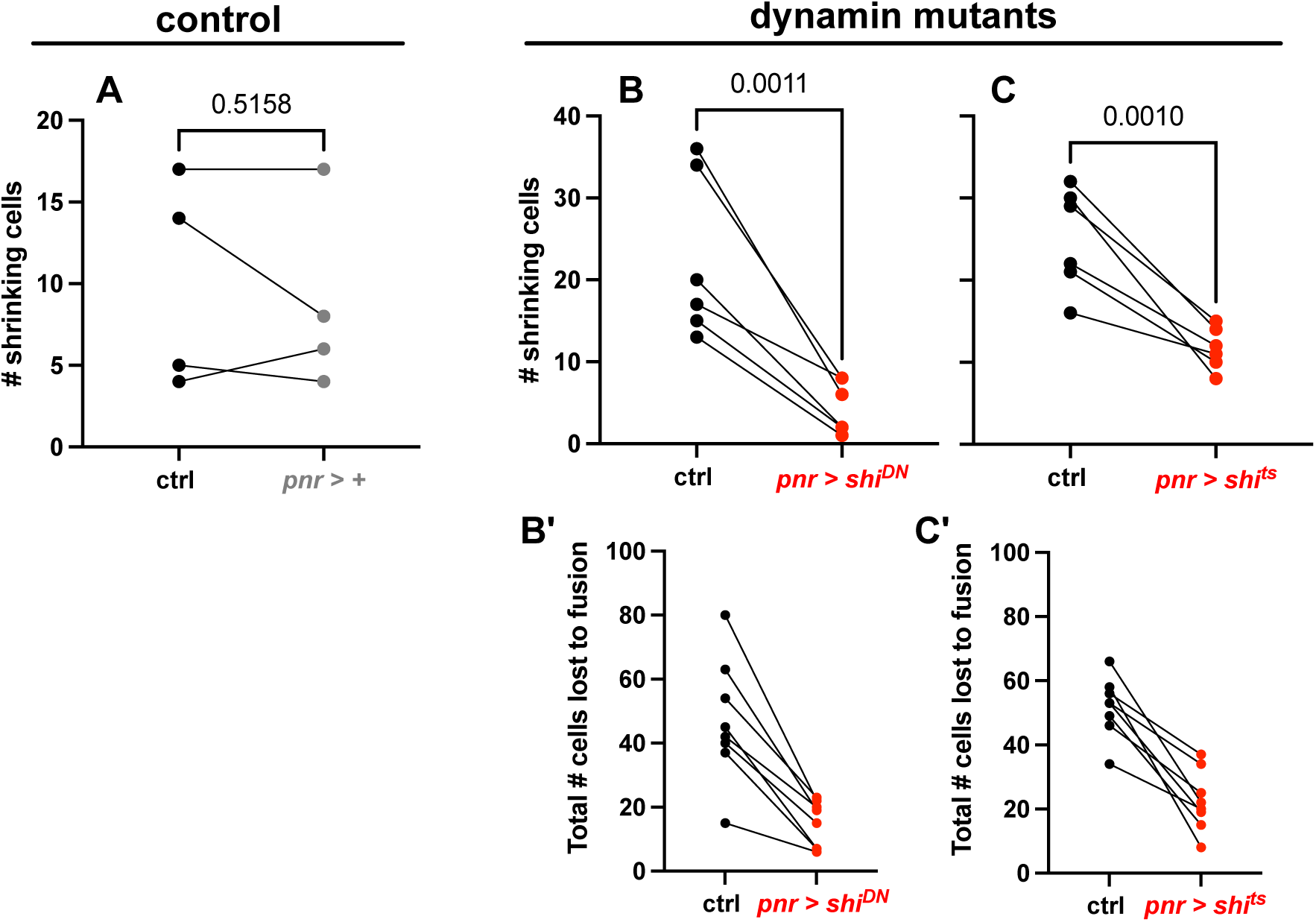
Wound-induced cell fusion can appear as the loss of adherens junctions as shown in Figures 1,2,5,6 or as cell shrinking as reported in White et al (2023) and analyzed here. **A.** In control pupae, the number of wound-induced shrinking-cell fusions is comparable on both sides of the wound (*pnr* and ctrl domains; n = 4 pupae; paired T test). **B-C.** Inhibition of shibire (dynamin) reduces the number of wound-induced shrinking-cell fusion events (B-C). Combining both fusion routes (B’-C’), inhibition via *shi^DN^* (n = 6 pupae) has an overall higher impact on wound-induced fusion than does *shi^ts^* (n = 6 pupae; paired T test). The number of shrinking cells was counted manually by tracking the Ecad signal throughout 5-hr movies.

**Figure S3:**
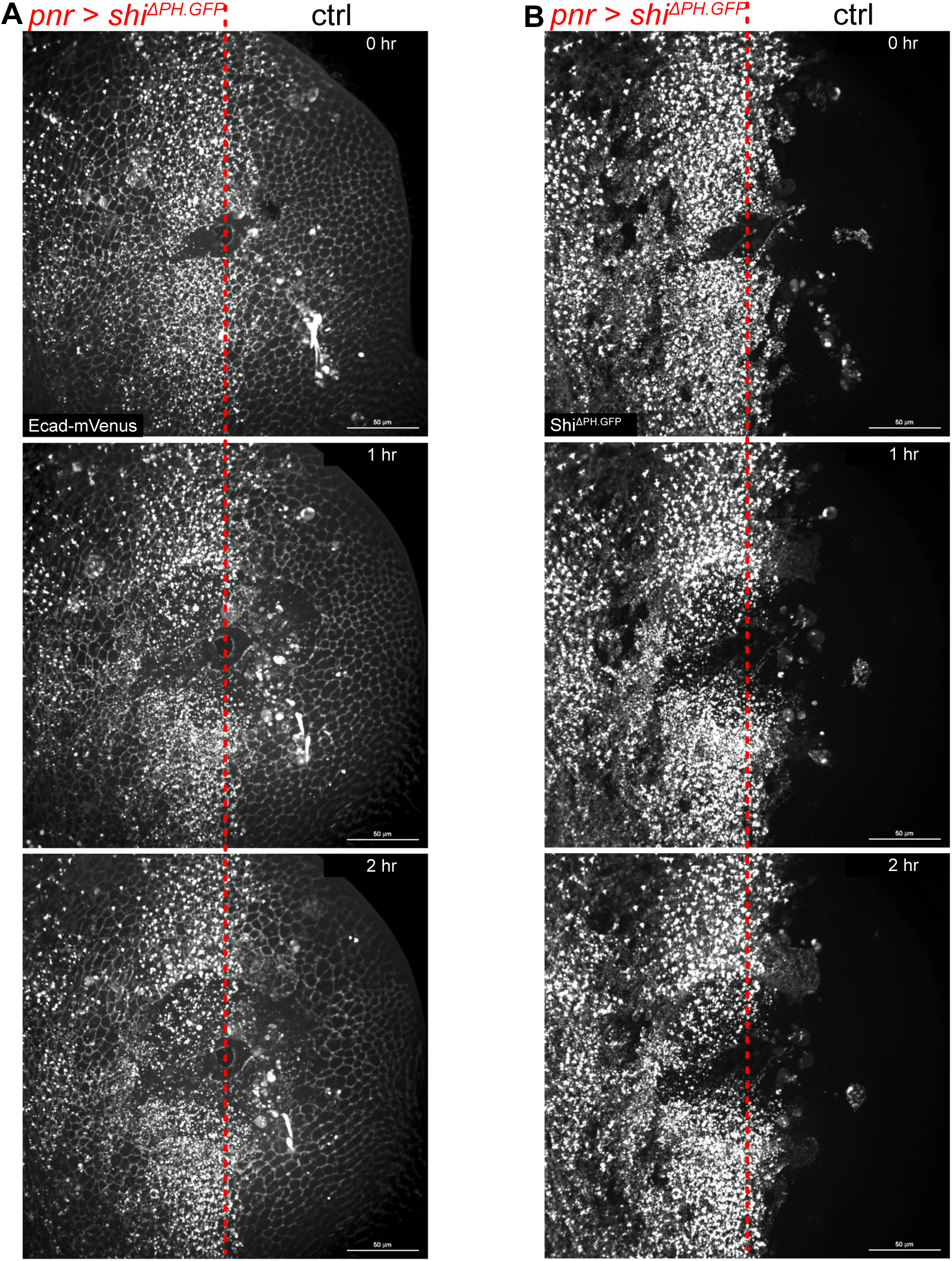
*shi^ΔPH^-GFP*, an endocytosis-specific *shi* mutant that cannot oligomerize around lipid nanotubes, impairs wound-induced fusion. **A, B.** *shi^ΔPH^-GFP* was expressed in the *pnr* domain. Ecad-mVenus labeling of cell borders is shown in A, and transgenic Shi^ΔPH^-GFP protein in the same wound is shown in B. Syncytia appear bigger in the control domain than in the *pnr* domain where *shi^ΔPH^-GFP* is expressed (n = 7 wounds). The accumulation of Ecad-mVenus in the *pnr* domain is a result of *shi^ΔPH^-GFP* impairing Ecad endocytosis. The frequency of fusion was not quantified due to the suboptimal quality of Ecad signal.

**Fig. S4:**
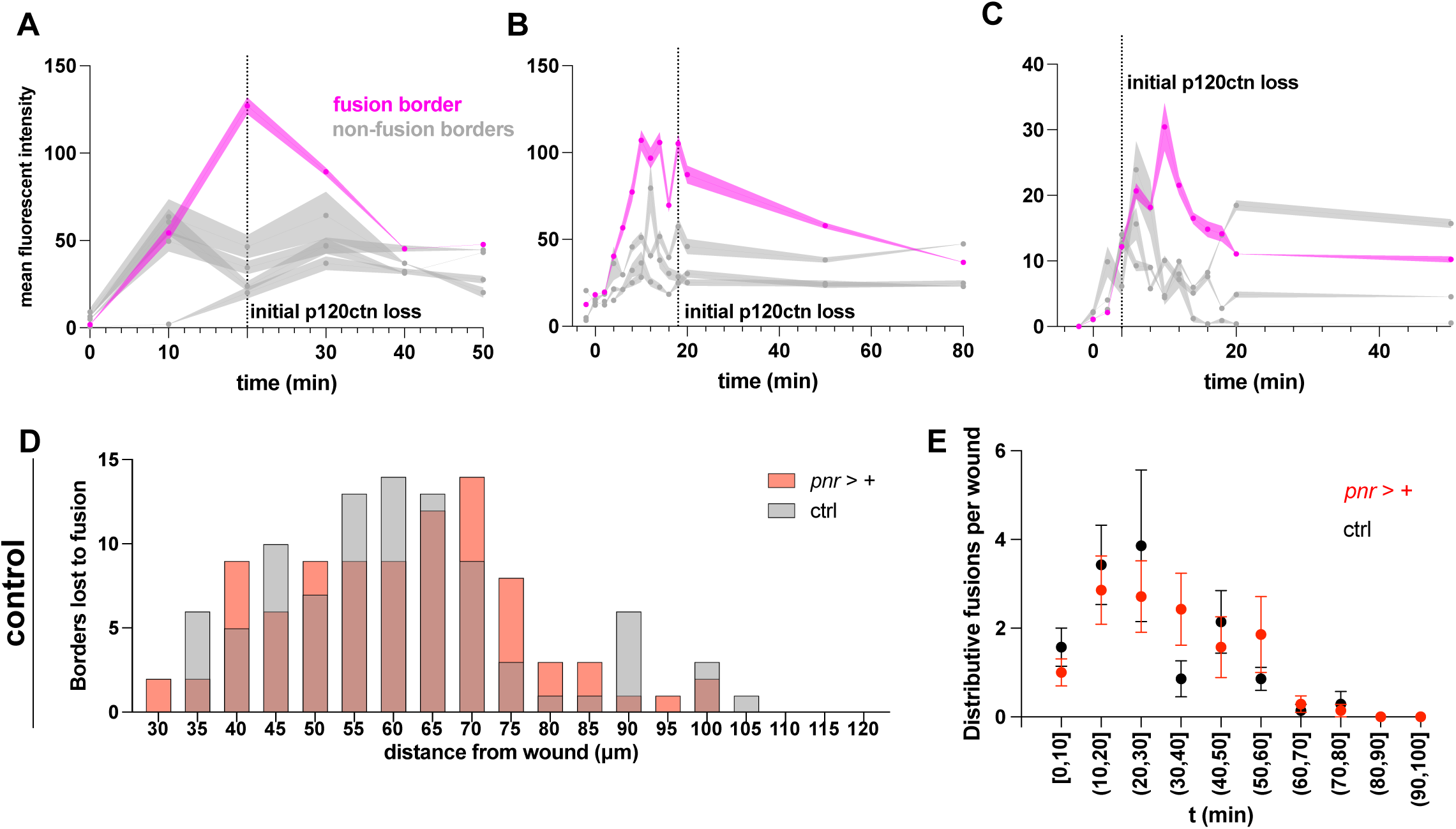
Spatiotemporal characterization of dynamin localization and function during wound-induced fusion. A-C. Three other examples of Shi-GFP localizing preferentially to the fusion cell border between 1° and 2° recipient cells after wounding. Line thickness represents SEM. D-D’ Both the spatial (D) and temporal (E) distribution of fusion is comparable in control and *pnr* domains without genetic manipulation (n = 7 wounds).

**Fig. S5:**
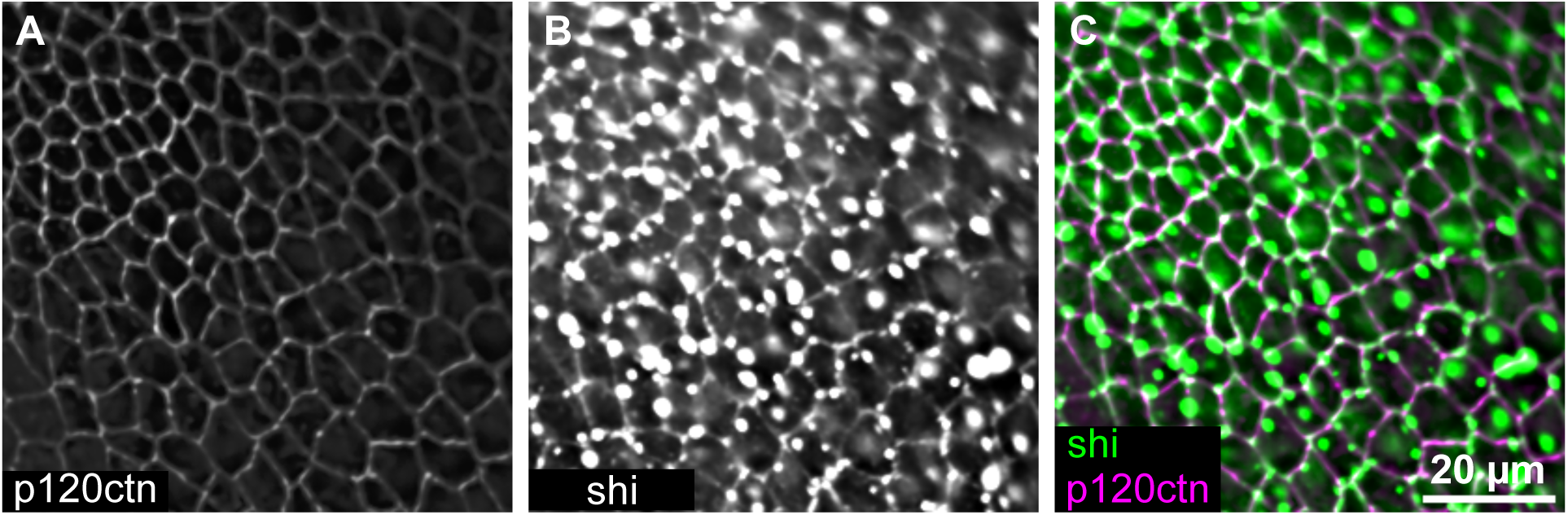
Expressing *shi-GFP* in all cells in the *pnr* domain generates a strong signal that obscures wound-induced changes in Shi-GFP localization. A-C. Single Z slice image of p120ctn (A) shi-GFP (B) and the merged image (C).

**Fig. S6:**
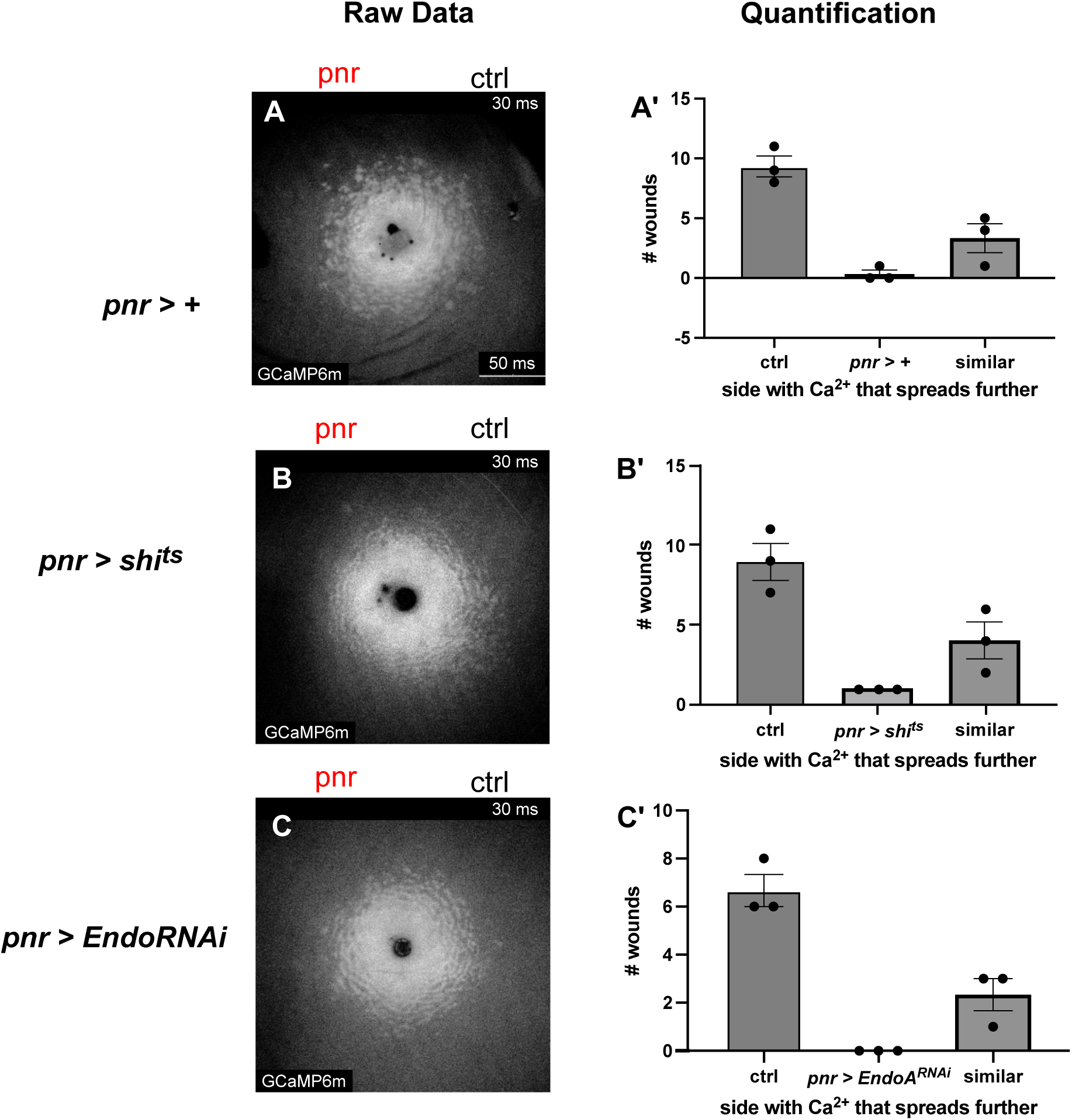
The extent of plasma membrane damage is not affected by the loss of dynamin or endophilin. We evaluated whether the level of wound-induced plasma membrane damage was altered when endocytosis was inhibited (endophilin or dynamin inhibition), as one potential explanation for the reduction in wound-induced cell fusion could be a reduction in the level of plasma membrane damage. We found no evidence that plasma membrane damage was altered in the absence of endocytosis. A-C. Calcium signatures imaged 30 ms after wounding with GCaMP6m appear comparable with and without functional *shi* or *EndoA*. A’-C’ After inhibiting either *shi* or *EndoA* (B’, C’), the inherent difference in calcium range between pnr and ctrl domains (A’) is not altered, indicating that the function of the two proteins does not affect the level of plasma membrane damage. Method: To compare the extent of calcium signal between *pnr* and control domains, three blinded observers were asked to describe shuffled 30 ms calcium images (38 total images: 13 control, 14 *shi^ts^*, 9 *EndoA-RNAi*) in terms of the extent of calcium imaging: did calcium spread further on the left side, right side, or were the two sides comparable in a given image? Responses were similar for all three genotypes.

**Fig. S7:**
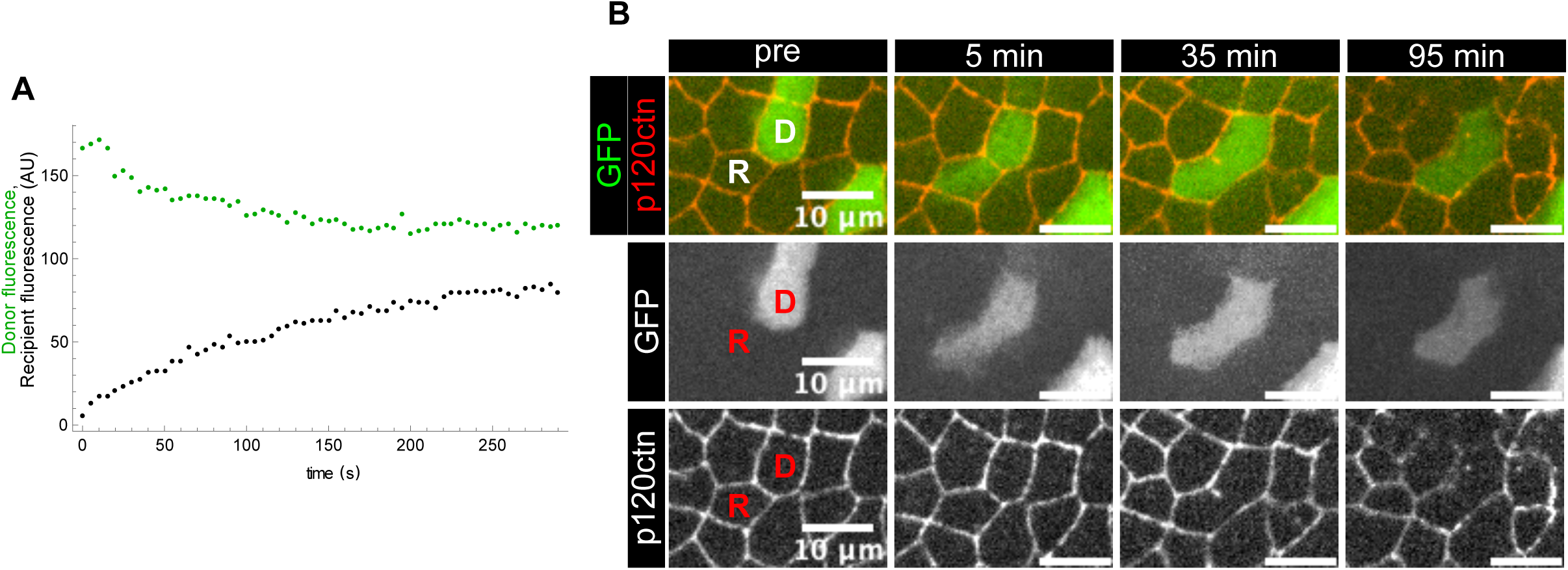
In control cell fusion events, even when cytoplasmic transfer becomes stalled, fusion occurs eventually. A. Imaging of cytoplasmic GFP within 5 min after wounding indicates that the flow of GFP from this donor cell (green dots) to recipient cell (black dots) became stalled, with decreasing permeance over time. B. Live imaging of the same donor and recipient over the next 1.5 h shows that eventually, the GFP equilibrated and the border between the two cells broke down, indicating that equilibration was reached and fusion was completed. Among the 19 pairs of donor-recipient cells analyzed in 6 pupae, all 10 pairs that did not reach equilibration within 5 min eventually underwent border breakdown and equilibrated.

**Figure S8:**
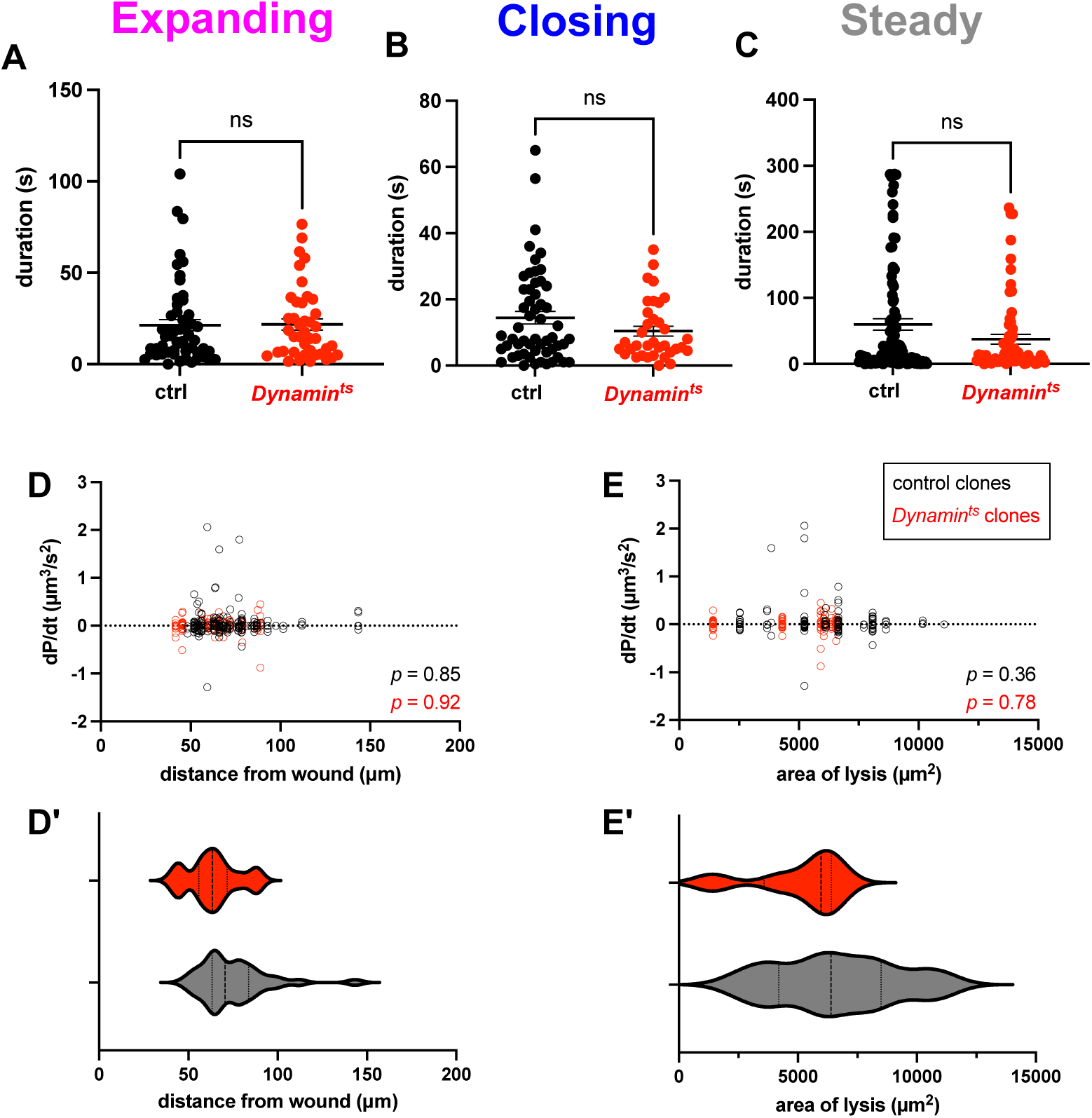
Further characterization of wound-induced cell-cell GFP permeance. A-C. Time spent at each of the three stages is comparable between control and *shi^ts^* clones (expanding: *p* = 0.96; closing: *p* = 0.42; steady: 0.50; *p* value calculated from Mann-Whitney test, error bar = 1 +/- SEM). D-D’. dP/dt, a readout for the speed of open channel expansion/closing, is not correlated with distance from wound in either the control clones or the *shi^ts^* clones. E-E’. dP/dt is not correlated with the size of the wound (area of lysis) in either the control clones or the *shi^ts^* clones. *p* values calculated from Pearson’s correlation test.

